# Correlations between stochastic endemic infection in multiple interacting subpopulations

**DOI:** 10.1101/568667

**Authors:** Sophie R Meakin, Matt J Keeling

## Abstract

Heterogeneity plays an important role in the emergence, persistence and control of infectious diseases. Metapopulation models are often used to describe spatial heterogeneity, and the transition from random-to heterogeneous-mixing is made by incorporating the interaction, or coupling, within and between subpopulations. However, such couplings are difficult to measure explicitly; instead, their action through the correlations between subpopulations is often all that can be observed. We use moment-closure methods to investigate how the coupling and resulting correlation are related, considering systems of multiple identical interacting populations on highly symmetric complex networks: the complete network, the *k*-regular tree network, and the star network. We show that the correlation between the prevalence of infection takes a relatively simple form and can be written in terms of the coupling, network parameters and epidemiological parameters only. These results provide insight into the effect of metapopulation network structure on endemic disease dynamics, and suggest that detailed case-reporting data alone may be sufficient to infer the strength of between population interaction and hence lead to more accurate mathematical descriptions of infectious disease behaviour.

## 1 Introduction

Heterogeneity is an increasingly important feature of epidemiological models, with spatial structure (Grenfell and Bolker, 1998; Xia et al., 2004; Viboud et al., 2006), risk structure (Baguelin et al., 2010; Datta et al., 2018; Rock et al., 2018) and age structure (Schenzle, 1984; Keeling and Grenfell, 1997; Keeling and White, 2010) widely considered. The incorporation of various forms of heterogeneity is crucial to capture many important observed epidemiological dynamics, such as: clustering of cases, either spatially or in high-risk demographics (Schenzle, 1984); unexpected endemic behaviour, as heterogeneity breaks down the simple formulation of the basic reproduction number (Keeling and Rohani, 2008); and persistence, where heterogeneity generally acts to increase persistence (Keeling, 2000; Hagenaars et al., 2004). Heterogeneity also has a marked influence on the control of infectious diseases, as a result of increased persistence or driven by targeted interventions (Keeling and White, 2010; Christley et al., 2005; Wallinga et al., 2010).

One modelling framework that can capture multiple forms of heterogeneity is the metapopulation-type model (Gilpin and Hanski, 1991; Hanski, 1998; Hanski and Gaggiotti, 2004), whereby the population is divided into multiple interacting, or ‘coupled’, subpopulations, and where within-population interactions typically occur at a higher rate than between-population interactions. Metapopulation models usually describe spatially distributed communities, but could also represent risk groups (e.g. high and low risk) or age groups (e.g. adults and children).

Quantifying between-population interactions is one of the key challenges of metapopulation infectious disease modelling (Ball et al., 2014). The individual-level behaviour that determines such interactions is highly complex and is dependent on social, cultural, environmental and economic factors (Wesolowski et al., 2015). Even with access to good data on relevant interactions, it is unclear how this should translate into a single phenomenological transmission parameter; current approaches to spatially structured metapopulation models typically combine theoretical models of human mobility with highly detailed data. Models of human mobility characterise the distribution of contacts between populations based on the population sizes and the distances between them (Hanski, 1998). The gravity model, originally formulated for transportation analysis (Erlander and Stewart, 1990), and later modified for infectious disease modelling, has been widely used in combination with commuter mobility data (Viboud et al., 2006; Balcan et al., 2009), mobile phone data, used as a proxy for human mobility (Tizzoni et al., 2014; Wesolowski et al., 2015; Kraemer et al., 2016), or spatiotemporal time series of disease incidence, where coupling parameters are estimated so that simulated epidemics match observed case numbers (Xia et al., 2004). However, good data on relevant movements between populations are rare, particularly in developing countries where epidemiological models are more likely to be applied. The parameter-free radiation model (Simini et al., 2012) and variants thereof (Yan et al., 2014; Kang et al., 2015) offer alternative models for human mobility that only requires the spatial distribution of the population to estimate coupling. However, comparisons between both the gravity and radiation models, and mobile call data records show that these models fail to fully describe human mobility outside of high-income countries, such as in Sub-Saharan Africa (Wesolowski et al., 2015).

The interaction between subpopulations is often represented as a matrix of transmission rates, where diagonal elements represent within-population rates and off-diagonal elements represent between-population rates. When considering *P* populations, this matrix has *P*^2^ elements, which leads to unidentifiability problems if attempting to estimate parameters from endemic equilibria. On the other hand, in a stochastic system, the 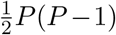 pairwise correlations between the levels of infection in pairs of populations may help to mitigate this unidentifiability, particularly if the transmission matrix is sparse or can be assumed to have some sort of symmetry. Long-term data on disease incidence is more frequently available (Olsen and Schaffer, 1990; Grenfell and Harwood, 1997), from which we can estimate the correlation between epidemics in distinct subpopulations; then, given an analytic relationship between the coupling and the correlation, we can infer interaction parameters.

Whilst computer simulations are commonly used and clearly useful, analytic results allow us to develop intuition about the infection dynamics; however, exact analysis of stochastic epidemiological models is often mathematically intractable, due to the nonlinearity of the transmission process. In such cases, approximation methods may be used to derive results about the expected behaviour and variability of the infection process. One such approximation method is a moment closure approximation, whereby the stochastic system is approximated by a deterministic system of differential equations for the moments (mean, variance, covariance, etc.). The most commonly used moment closure approximation, and the one used throughout this paper, is the multivariate normal approximation, which assumes that third-order cumulants and higher are equal to zero or, equivalently, that the distribution of states follows a multivariate normal (MVN) distribution (Whittle, 1957).

In this paper we derive an approximation for the correlation between the level of infection in two subpopulations as a function of the coupling between them. Our results extend the analysis of Meakin and Keeling (2018) for a simple two subpopulation system. Using a multivariate normal approximation we derive results for subpopulations arranged on the complete network, the *k*-regular tree network and the star network. We also numerically validate our model by comparing our analytic approximations to stochastic simulations. These results also provide some insight into the effect of metapopulation network structure on network correlations.

## 2 Methods

### 2.1 A stochastic endemic infection model for interacting populations on a general graph

We begin by introducing a simple stochastic *SIR* model in a population of size *N*, with births, deaths, transmission and recovery. At any time *t* ∈ [0, ∞), individuals are in one of three states: susceptible, infected or recovered. A given susceptible individual meets other individuals at rate *m* > 0. We assume that these encounters are sufficiently close that if the other individual is infected, then transmission of infection occurs with probability *τ* and the susceptible individual immediately becomes infected and infectious to others. We therefore define the transmission rate be *β* = *mτ*. Susceptible individuals can also succumb to infection independent of contact with infected individuals in the modelled populations; this occurs at rate *ϵ* > 0, the external import rate. Infected individuals recover from infection at rate *γ* > 0, after which they become immune to further infection. Susceptible, infected and recovered individuals all die at rate *μ* > 0, independent of infection status; we assume that a death is immediately followed by the birth of a susceptible individual, and hence the total population size remains constant. The basic reproductive ratio, the expected number of secondary cases produced by a single infectious individual in a susceptible population, for this process is *R*_0_ = *β*/(*γ* + *μ*).

We extend the above model to *P* identical populations of size *N*. The assumption that the population sizes are equal is for mathematical tractability; a discussion of the effects of relaxing this assumption for *P* = 2 can be found in Meakin and Keeling (2018). Each population exhibits the same population dynamics as described above, plus pairwise interaction between the populations: we assume that in population *i*, a proportion *σ_ij_* ∈ [0,1] of an individual’s contacts are with individuals in population *j*. We insist that 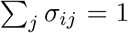 and so 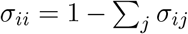. The matrix **Σ** = (*σ_ij_*) therefore describes the interaction or ’coupling’ between all possible pairs of populations, and the force of infection in each subpopulation depends on the number of infected individuals in all other subpopulations. Changing **Σ** does not change the basic reproductive ratio, but instead determines the distribution of secondary cases between the *P* subpopulations.

We let *S_i_*(*t*), *I_i_*(*t*), *R_i_*(*t*) ∈ {0,1,2,…} denote the number of susceptible, infected and recovered individuals, respectively, in population *i* = 1, 2,…, *P* at time *t* ≥ 0. As the population size N is constant then *S_i_*(*t*)+ *I_i_*(*t*)+ *R_i_*(*t*) = *N*, ∀*t* ≥ 0, *i* = 1, 2,…, *P*. The transition rates for the resulting 2*P*-dimensional Markov chain from state (*s*_1_, *i*_1_, *s*_2_, *i*_2_,…, *s_P_*, *i_P_*) at time *t* are summarised in Table 1.

**Table 1.**
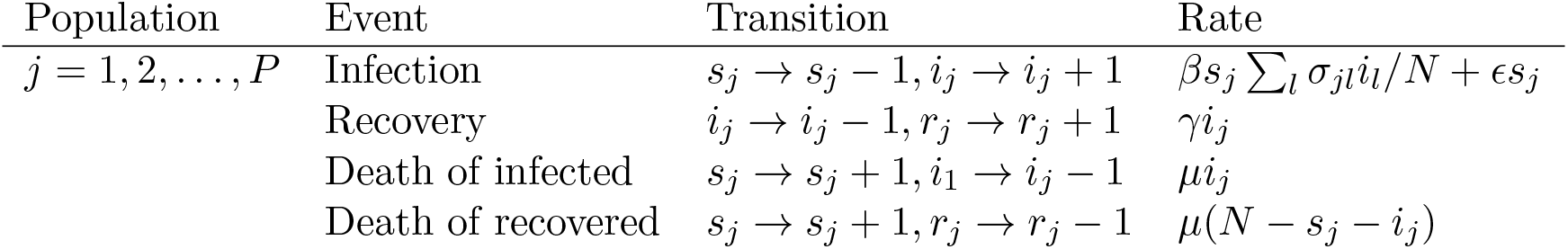
A summary of the transition rates of the 2*P*-dimensional Markov chain endemic infection model 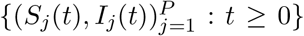 from state (*s*_1_, *i*_1_, *s*_2_, *i*_2_,…,*s_P_*, *i_P_*) with birth/death rate *μ* > 0, contact rate *β* > 0, external import rate *ϵ* > 0, recovery rate *γ* > 0 and coupling matrix **Σ**.

| Population | Event | Transition | Rate |
| --- | --- | --- | --- |
| $j = 1, 2, \dots, P$ | Infection | $s_j \rightarrow s_j - 1, i_j \rightarrow i_j + 1$ | $\beta s_j \sum_l \sigma_{jl} i_l / N + \epsilon s_j$ |
| | Recovery | $i_j \rightarrow i_j - 1, r_j \rightarrow r_j + 1$ | $\gamma i_j$ |
| | Death of infected | $s_j \rightarrow s_j + 1, i_1 \rightarrow i_j - 1$ | $\mu i_j$ |
| | Death of recovered | $s_j \rightarrow s_j + 1, r_j \rightarrow r_j - 1$ | $\mu(N - s_j - i_j)$ |

The metapopulation structure can be described by a weighted network *G* = (*V, E*) with vertex set *V* = {1,2,…, *P*} and edge set *E*, where edge *e* = *ij* has weight *σ_ij_*: the coupling matrix **Σ** therefore represents the weighted adjacency matrix for the graph *G*. For mathematical tractability we restrict our analysis to networks for which we can derive analytic results, namely graphs that are highly symmetric; a discussion of the effect of relaxing this assumption is provided in the Supplementary Information. In the following analysis we consider the complete network, the *k*-regular tree network and the star network. In addition, we assume that *σ_ij_* = *σ*, ∀*ij* ∈ *E*. We note that for *k*-regular tree network and the star network, the weighted adjacency matrix **Σ** is sparse, that is, most of the elements are zero.

### 2.2 Moment closure approximations

Even with constraints on the metapopulation network structure and the coupling matrix **Σ**, an exact analysis of the full stochastic model is mathematically intractable. Instead we consider the approximate behaviour of the first- and second-order central moments of the process. The ODE for 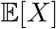 can be calculated from first principles using:

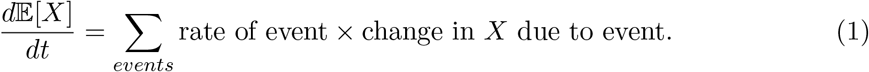

Alternatively, these ODEs can be derived from the Kolmogorov forward equation; details of this method can be found in existing literature on moment closure approximations in infectious disease modelling (Keeling and Rohani, 2002; Lloyd, 2004).

Due to the nonlinearity of the infection term in the model, the ODE for an *n*th-order moment will depend on one or more (*n* + 1)th order moments: to fully describe the stochastic process would therefore require an infinite set of ODEs. To circumvent this problem, we use a moment closure approximation, which truncates the set of ODEs at some order. Throughout this paper, we use a second-order moment closure approximation, which assumes that third- and higher-order cumulants are equal to zero. As a result, third- and higher-order moments can be written in terms of the means, variances and covariances only.

Throughout this paper we will use the following notation for the first- and second-order central moments:

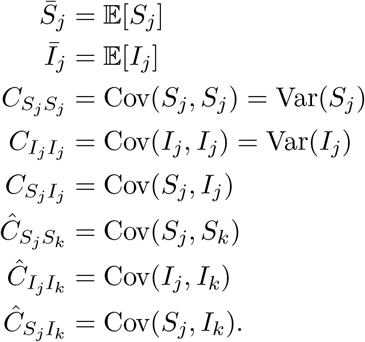

For a metapopulation network on *P* populations, the set of ODEs approximating the stochastic process has at most 3*P*^2^ + 2*P* equations: *P* for each of the two first order moments and *P*^2^ for each of the three covariances. However, for the networks that we consider in this paper, symmetries in the structure of the network mean that the number of ODEs is considerably fewer. In some cases we will simplify the notation: we outline simplifications to the notation at the start of the results section for each network.

### 2.3 Deriving an equation for the correlation

In each metapopulation network (the complete network, the *k*-regular tree network and the star network), we derive an analytic approximation for the correlation between the number of infected individuals in a pair of populations as a function of the coupling *σ*. We define the correlation between the number of infected individuals in population *i* and the number of infected individuals in population *j* at endemic equilibrium as:

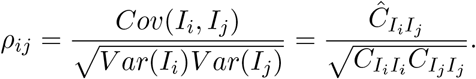

We derive an approximate equation for the correlation *ρ_ij_* by considering the ODE for the covariance 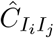 at endemic equilibrium. We then evaluate our approximation numerically, for which we need to define a set of base parameters. We utilise parameters for a highly-transmissible measles-like endemic disease in the UK (Anderson and May, 1992), although we note that a full model of measles requires both seasonality (Earn et al., 2000; Rohani et al., 2002; Grenfell and Bolker, 1995) and age-structure (Schenzle, 1984; Keeling and Grenfell, 1997; Bolker, 1993). We consider the effect of both the coupling and other parameters on the correlation; we also evaluate the accuracy of our approximation by comparing our results to simulations.

## 3 Results

### 3.1 The complete network

#### 3.1.1 Network definition and notation

First we consider *P* identical populations on the complete network, where each population interacts with the other *k* = *P* − 1 populations: a visual representation of the complete network for *P* = 3 and *P* = 5 populations is given in Figure 1. The coupling matrix **Σ** = (*σ_ij_*) is defined as

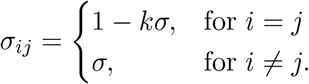

**Figure 1.**
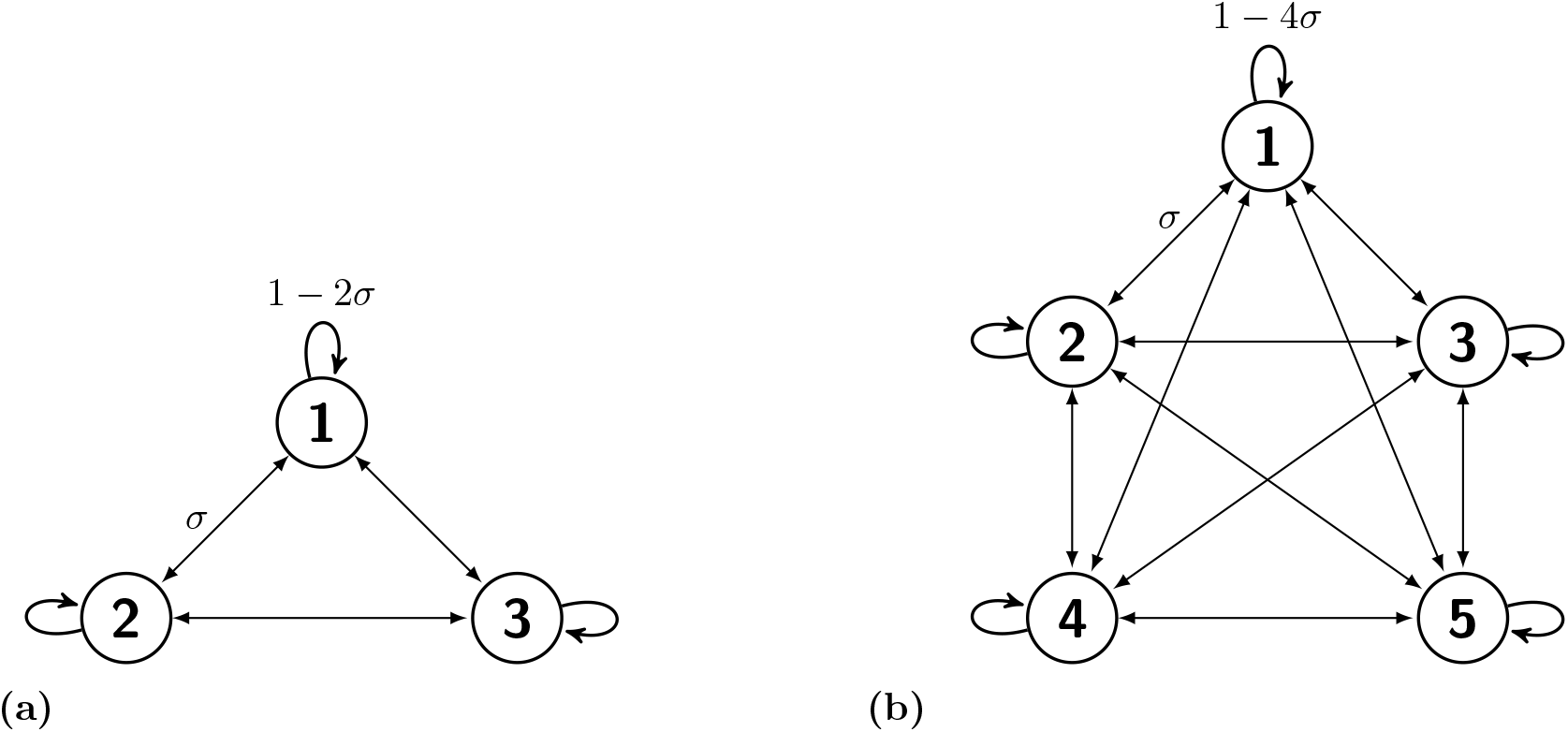
The complete network on (a) *P* = 3 and (b) *P* = 5 populations. The coupling between any pair of populations coupling is *σ* ∈ [0,1/(*P* − 1)] and so the within-population coupling is 1 − (*P* − 1)*σ*.

In the complete network metapopulation all subpopulations are epidemiologically and spatially identical: epidemiologically in the sense that all subpopulations are of equal size and have identical epidemiological parameters, and spatially in the sense that all nodes are isomorphic within the network and the coupling is the same between any pair of subpopulations. As a result, the expected behaviour is the same within all populations, and between any pair of populations. In our notation, we can therefore drop dependency on the population and simplify it to the following: 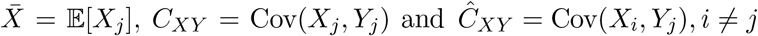.

Using the second-order moment closure approximation, and with these simplifications, the stochastic process on the complete network can be approximated by a set of eight ODEs: five for the within-population moments, and three for the between-population moments. These can be found in the Supplementary Information. We use these equations in both the analytic and the numerical results.

#### 3.1.2 Analytic results

For *P* populations on the complete network, we define the correlation between any pair of populations as

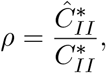

and show that this is equal to

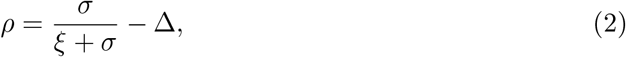

where

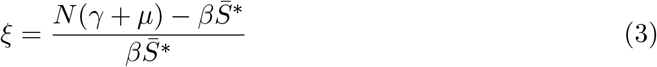

and

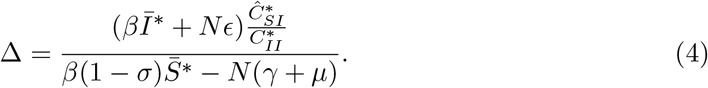

We derive this result by taking the moment equation for 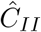 at equilibrium and dividing through by 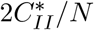, following the same approach as Meakin and Keeling (2018); full details of this derivation can be found in the Supplementary Information. Moreover, if Δ ≪ 1 then we can further simplify the approximation for the correlation to the following expression:

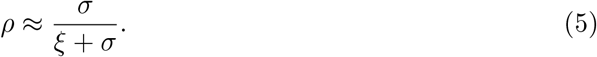

We can also use an alternative approximate expression for *ξ* that is independent of 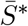, which eliminates the need to find the equilibrium of the 8-dimensional ODE model. Meakin and Keeling (2018) show that by ignoring the effects of imports and correlations and taking the large population limit, then

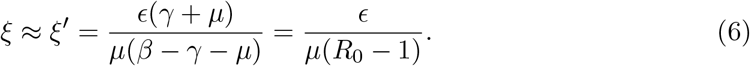

Given the simpler form of Equation (6) compared to the original expression for *ξ* given by Equation (3), in remainder of the analysis we evaluate *σ*/(*ξ^′^* + *σ*) as an approximation for the MVN correlation ρ.

This approximation is independent of the number of populations P. In short, this is due to the balance between two competing influences: the addition of an extra external coupling would normally weaken the correlation between two connected populations, but the fact that this additional population is itself correlated with the original populations nullifies this effect. In the Supplementary Information, we make this argument explicit by adding a third population (with variable coupling) to an interacting pair of populations.

#### 3.1.3 Numerical results

We first explore the effect of the number of subpopulations *P* and coupling *σ* on the equilibrium values of the first-order central moments 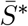 and 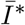 and the second-order central moments 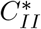 and 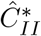 (Figure 2a). We consider *P* = 3, 5,10 and *σ* ∈ [0,1/*k*], *k* = *P* − 1, and include *P* = 2 for comparison. These results are obtained by the numerical integration of the system of ODEs given in the Supplementary Information, and so only introduce an error due to the MVN moment closure approximation. For all values of *P*, all curves show a sigmoidal pattern, with 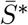 and 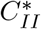 decreasing with the coupling, and 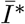 and 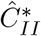 increasing with the coupling. As the number of populations *P* increases the magnitude of change in 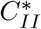 increases, since reducing the within-population coupling (either by increasing the between-population coupling *σ* or increasing the number of populations *P*) reduces the variance *C_II_*. However, the magnitude of change in 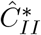 decreases, because as *P* increases, then the effect of interaction between a subpopulation and its neighbour is damped by the other *P* − 2 neighbours. In the previous section we noted that our approximation for the correlation is independent of the number of populations *P*: we also calculate the MVN correlation 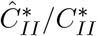 (Figure 2b) and note that this also appears independent of *P*. The correlation follows a sigmoidal relationship, increasing from zero for very low coupling.

**Figure 2.**
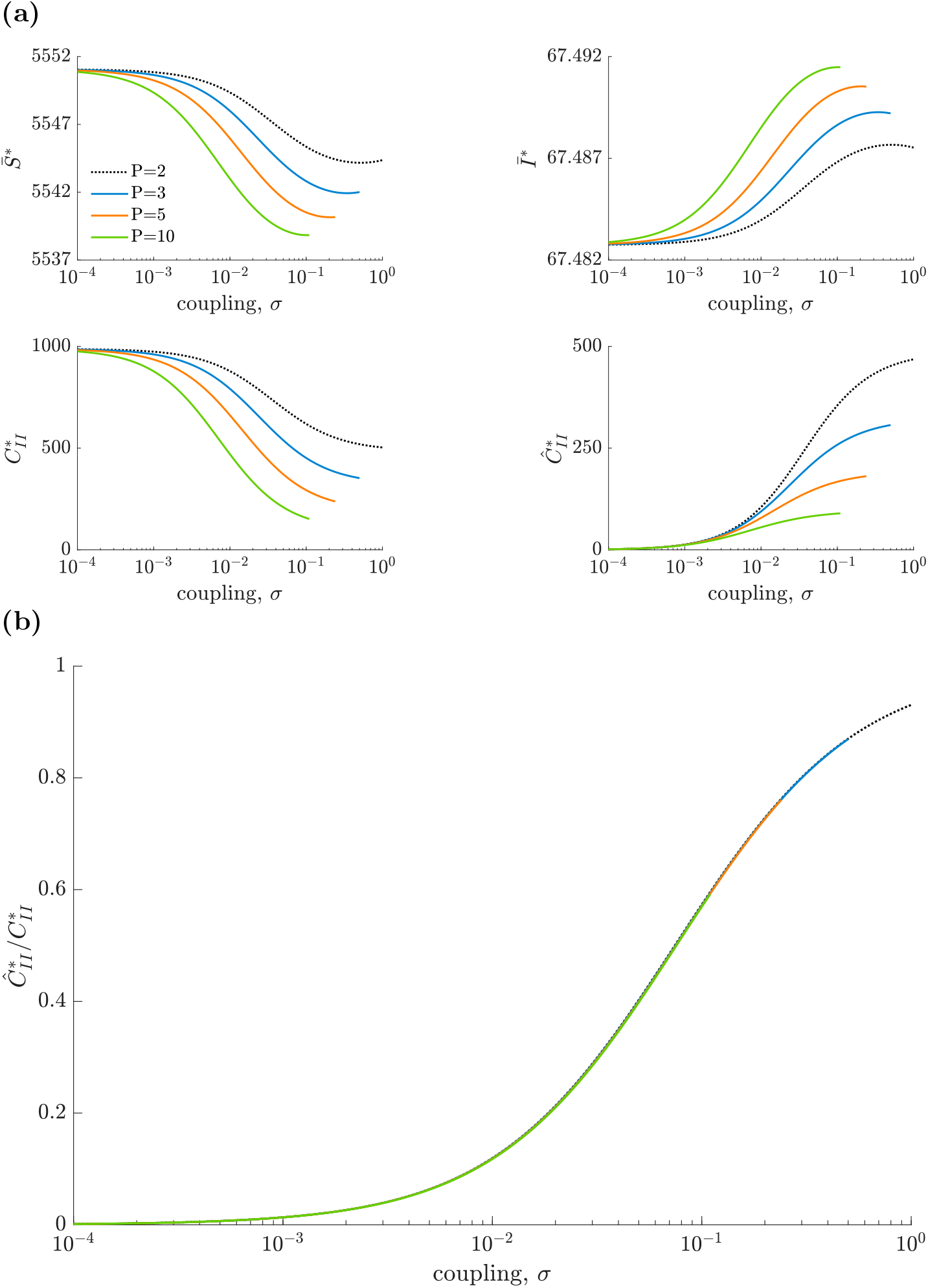
The effect of the coupling *σ* on (a) the key mean variables 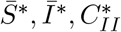 and 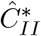 and (b) the correlation 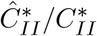, for *P* populations arranged on the complete network. Parameter values represent a measles-like endemic disease in the UK (*N* = 10^5^, *μ* = 5.5 × 10^−5^, *R*_0_ = 17, *γ*^−1^ = 13 and *ϵ* = 5.5 × 10^−5^). These values are calculated from the system of ODEs given in the Supplementary Information.

Next we compare the MVN correlation *ρ* (Equation (2)) and our simplified approximation *σ*/(*ξ^′^* + *σ*), *ξ^′^* = 0.0625 (Equation (5)) to stochastic simulations for *P* = 3, 5 subpopulations (Figure 3). The close agreement between *ρ* and the simulation results suggests that our use of the MVN moment closure approximation is justified. There is also little difference between the MVN correlation and our approximation (that is, Δ is small), so *σ*/(*ξ^′^* + σ) is a good approximation for the correlation *ρ*. Therefore, we can relate the phenomenological coupling parameter *σ* to the correlation between the number of infected individuals in any pair of populations for *P* populations arranged on the complete network by *ρ* ≈ *σ*/(*ξ^′^* + *σ*).

**Figure 3.**
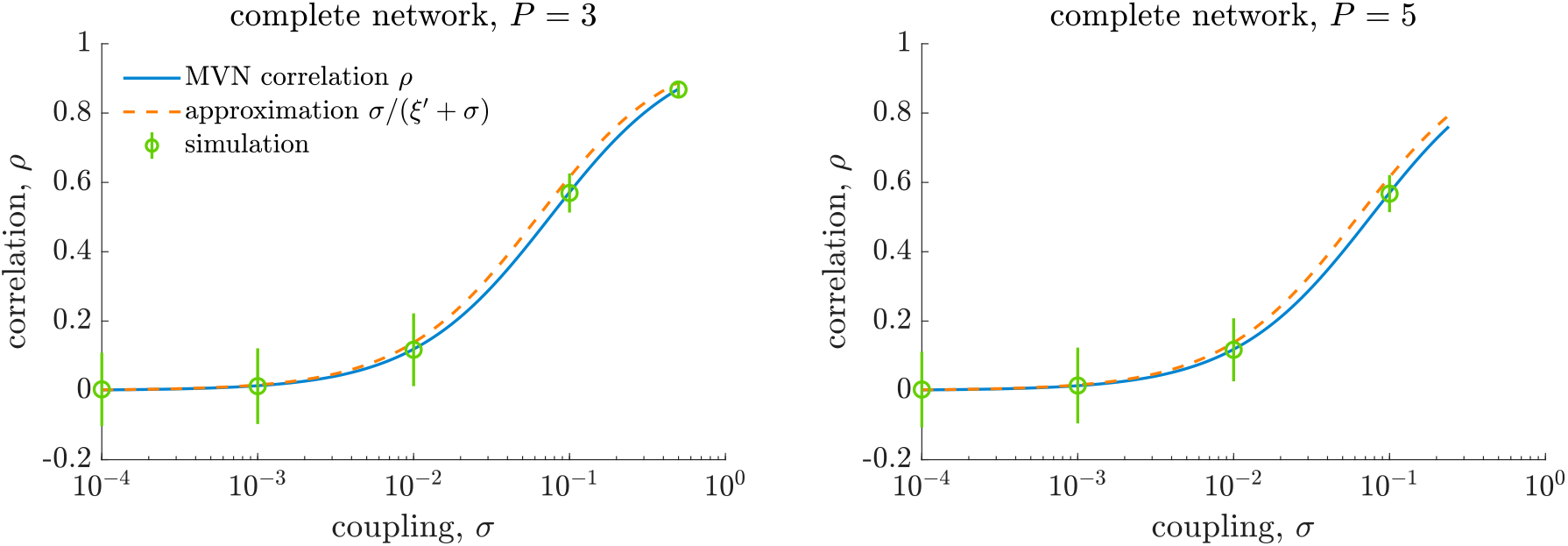
Comparing analytic and numerical correlation between any pair of populations from *P* = 3,5 populations arranged on the complete network. We compare the analytic correlation *ρ* and our approximation *σ*/(*ξ^′^* + *σ*),*ξ^′^* = 0.0625, to stochastic simulations for a measles-like endemic disease in the UK (*N* = 10^5^, *μ* = 5.5 × 10^−5^, *R*_0_ = 17, *γ*^−1^ = 13 and *ϵ* = 5.5 × 10^−5^). Each population is coupled to the *k* = *P* − 1 other populations. The between-population coupling is fixed as *σ* ∈ [0,1/*k*] and within-population coupling is therefore 1 − *kσ*. We generate 1000 realisations of the process for each value of *σ* and calculate the correlation as a time-weighted Pearson correlation coefficient for 50 ≤ *t* ≤ 200; error bars represent ±2 standard deviations.

### 3.2 The tree network

#### 3.2.1 Network definition and notation

Next, we consider infinitely many populations on a *k*-regular tree network, where each subpopulation has *k* neighbours: a visualisation of the *k*-regular tree network for *k* = 2 and *k* = 4 neighbours is given in Figure 4. The coupling matrix **Σ** = (*σ_ij_*) is defined as

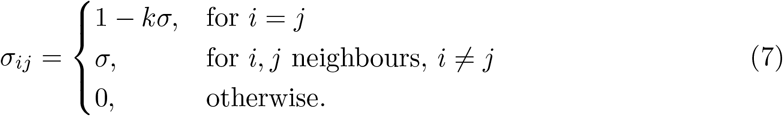

**Figure 4.**
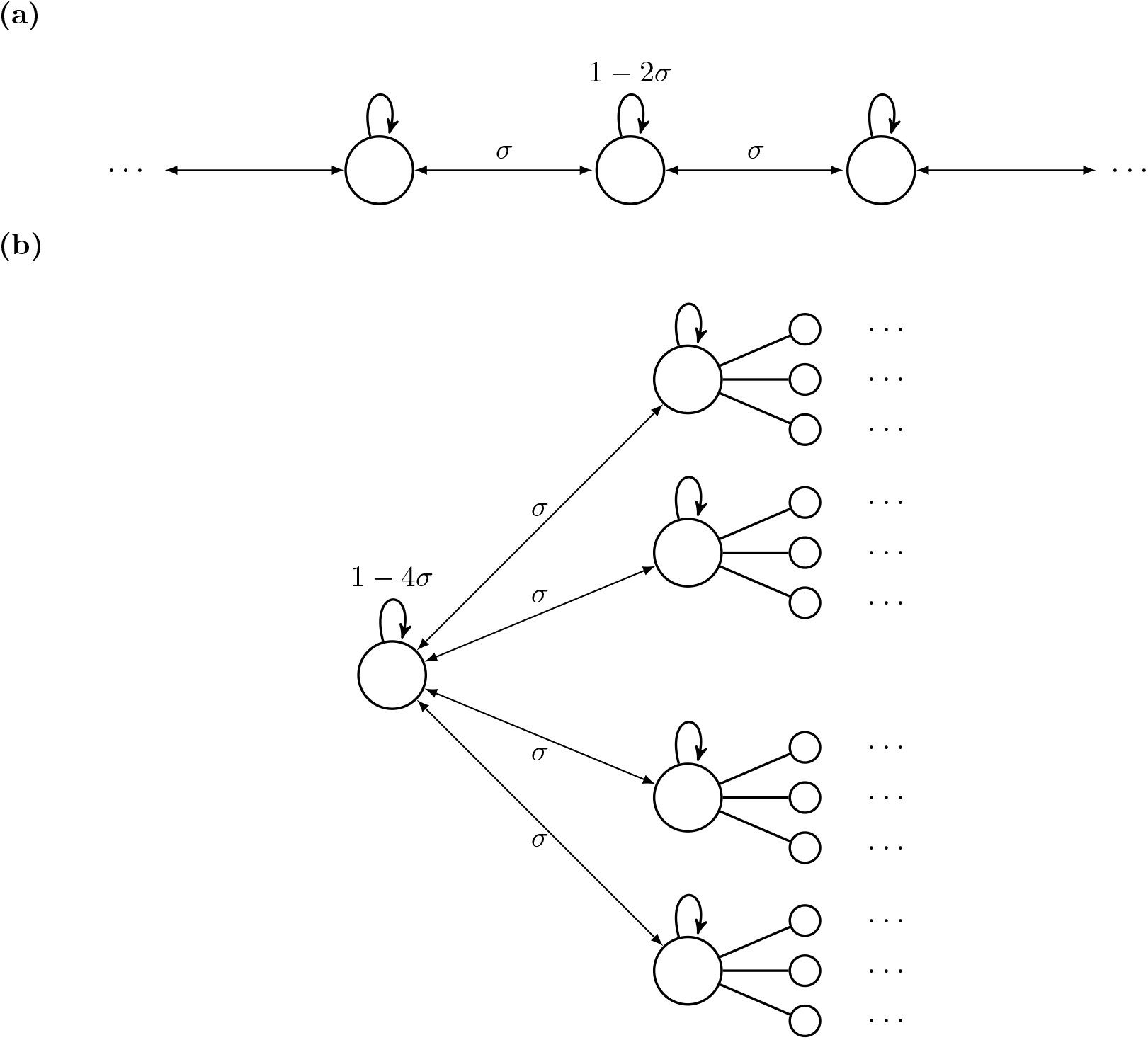
The *k*-regular tree network for (a) *k* = 2 and (b) *k* = 4 neighbours. The coupling between any pair of neighbouring populations is *σ* ∈ [0,1/*k*] and so the within-population coupling is 1 − *kσ*.

As with the complete network, all subpopulations in the *k*-regular tree network are epidemiologically and spatially identical, so the expected behaviour is the same within all subpopulations. In addition, in a tree network, there is a unique path between any pair of subpopulations, and so we can define the distance 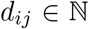 between subpopulations *i* and *j* to be the length of the path between the subpopulations. For the notation for within-population moments we can again drop dependency on the subpopulation: 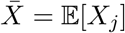 and *C_XY_* = Cov(*X_j_, Y_j_*). For the between-population moments, we only need to denote the distance *d* between the subpopulations: 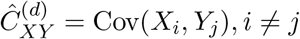, where *d_ij_* = *d*.

##### Finite subgraph approximation of the *k*-regular tree network

We cannot perform stochastic simulations of the infection process on infinitely many subpopulations. In addition, we can use a second-order moment closure approximation to derive a system of ODEs that approximate the stochastic process on the network, but this system comprises infinitely many equations: five equations for the within-population moments, and infinitely many equations for the between-population moments (3 for each *d* ≥ 1).

To overcome these problems, we consider a finite subgraph of the *k*-regular tree network. We define the *D*-truncated *k*-regular tree network to be the network of subpopulations distance less than or equal to *D* from some arbitrarily chosen origin node; since all subpopulations are identical and the *k*-regular tree network is infinite, the choice of origin node is irrelevant. The total number of subpopulations in the *D*-truncated *k*-regular tree network is

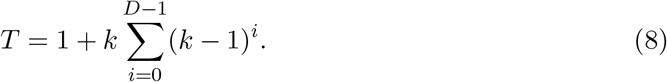

We can also write down a finite set of ODEs that approximate the stochastic process on the *D*-truncated *k*-regular tree network. If *D* is sufficiently large, then we can make some further simplifying assumptions. First, we can assume that 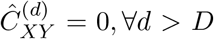. Secondly, we can assume that the expected behaviour of the first- and second-order central moments in the origin node, and between the origin node and subpopulations at distance *d* ≪ *D* will be the same as in the full *k*-regular tree network. In the full *k*-regular tree network we had that 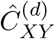 is the same for any pair of subpopulations distance *d* apart: we continue to make this simplification in the truncated network. Given these assumptions, and making a second-order MVN moment closure approximation, the stochastic process on the *D*-truncated *k*-regular tree network can be approximated by a set of 5 + 3*D* equations: five equations for the within-population moments and 3*D* equations for the between-population moments. These can be found in the Supplementary Information.

#### 3.2.2 Analytic results

We can derive analytic results for the full *k*-regular tree network. We define the correlation between the number of infected individuals in a pair of subpopulations distance *d* ≥ 1 apart as

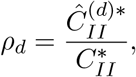

where *ρ*_0_ = 1 and 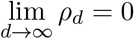. We can show that *ρ_d_* is the solution to

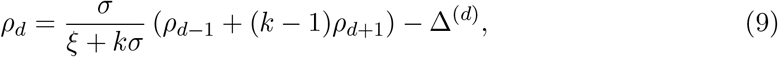

where

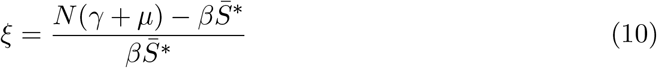

and

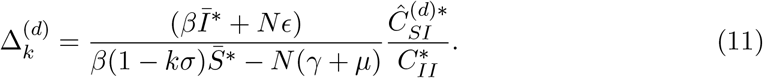

We derive this result from the moment equation for 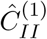 at equilibrium and dividing through by 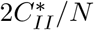; full details of this derivation can be found in the Supplementary Information. Moreover, if Δ^(*d*)^ ≪ 1, ∀*d* then *ρ_d_* is the solution to the recurrence relation

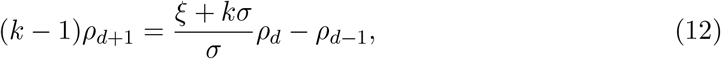

where *ρ*_0_ = 1 and 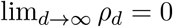. Since |*ρ_d_*| ≤ 1 then the solution is given by

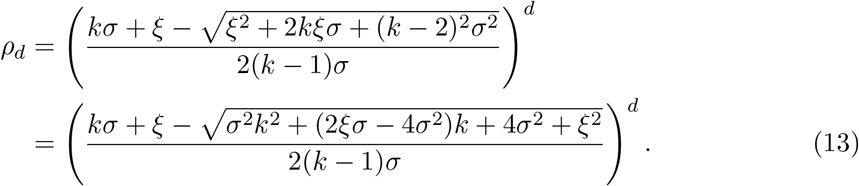

We note two things: firstly, since *ρ*_1_ ≤ 1 then it is trivial that *ρ_d_* → 0 as *d* → ∞. Secondly, *ρ_d_* → 0 as *k* → ∞.

#### 3.2.3 Numerical results

We note that the MVN correlation and stochastic simulations have to be performed on the *D*-truncated *k*-regular tree network, as it is not possible to use the full *k*-regular tree network. If *D* is sufficiently large, then these correlations will be approximately the same as in the full *k*-regular tree network: we show that for *D* sufficiently large then the correlation converges (Figure S2, Supplementary Information).

We first numerically evaluate the effect of the number of neighbouring subpopulations *k* and the distance *d* on the correlation *ρ_d_* (Figure 5). As with the complete network, the correlation follows a sigmoidal shape, increasing from zero correlation from very low coupling. For fixed coupling *σ*, as the number of neighbours *k* increases then the correlation *ρ_d_* decreases; similarly, for a fixed number of neighbours *k*, as the distance *d* increases then the correlation *ρ_d_* also decreases. This all agrees with expected behaviour from Equation (13).

**Figure 5.**
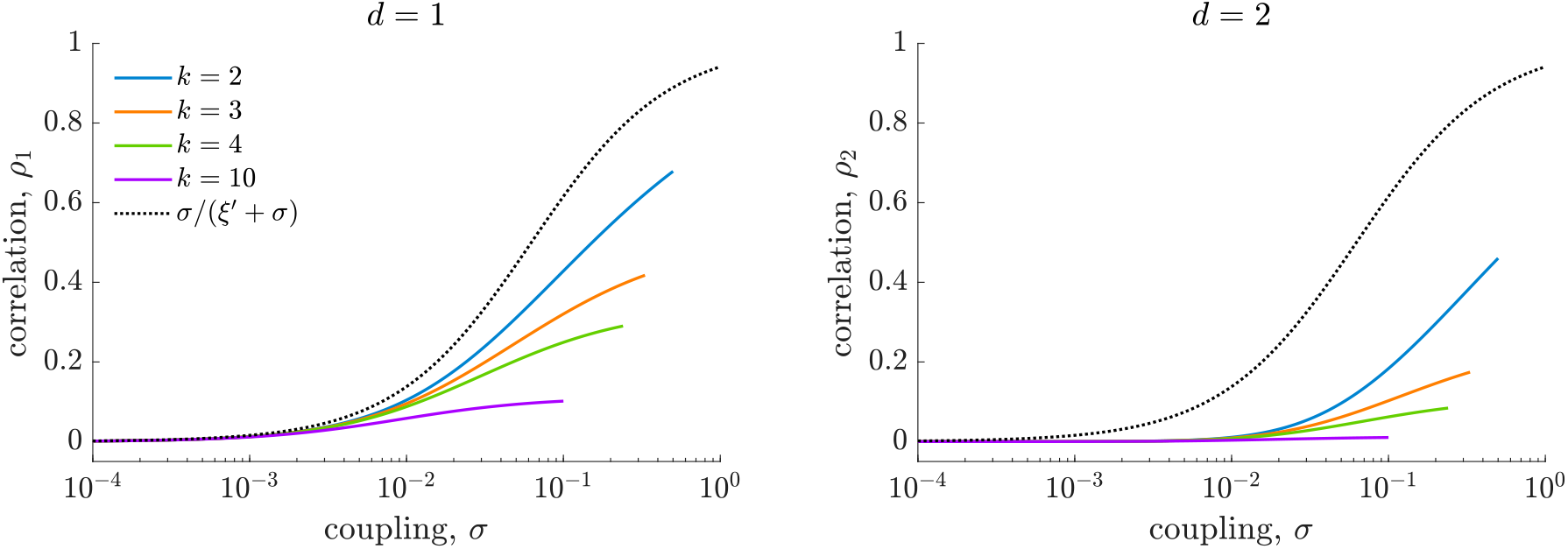
The effect of the number of neighbouring subpopulations *k* in the *k*-regular tree network on the correlation between the number of infected individuals in adjacent populations, *ρ*_1_ (left), and populations with a common neighbour, *ρ*_2_ (right). Parameter values represent a measles-like endemic disease in the UK (*N* = 10^5^, *μ* = 5.5 × 10^−5^, *R*_0_ = 17, *ϵ* = 5.5 × 10^−5^, *γ* = 1/13). The MVN correlation is calculated on the *D*-truncated *k*-regular tree network for *D* = 50 from the system of ODEs given in the Supplementary Information.

Next, we compare our approximations to the results of stochastic simulations for *k* = 2,4 (Figure 6), where stochastic simulations are performed on the *D*-truncated *k*-regular tree network and *D* = 5, 3 for *k* = 2, 4, respectively. For all combinations of *k* and *d* there is close agreement between the MVN correlation and stochastic simulations, which justifies our use of the MVN moment closure approximation; we can show that increasing *D* further does not significantly change the correlations in the system (Supplementary Information, Figure S2). There is also little difference between the MVN correlation and our approximation (that is, Δ^(1)^ is small) and so approximating the MVN correlation by Equation (13) is reasonable.

**Figure 6.**
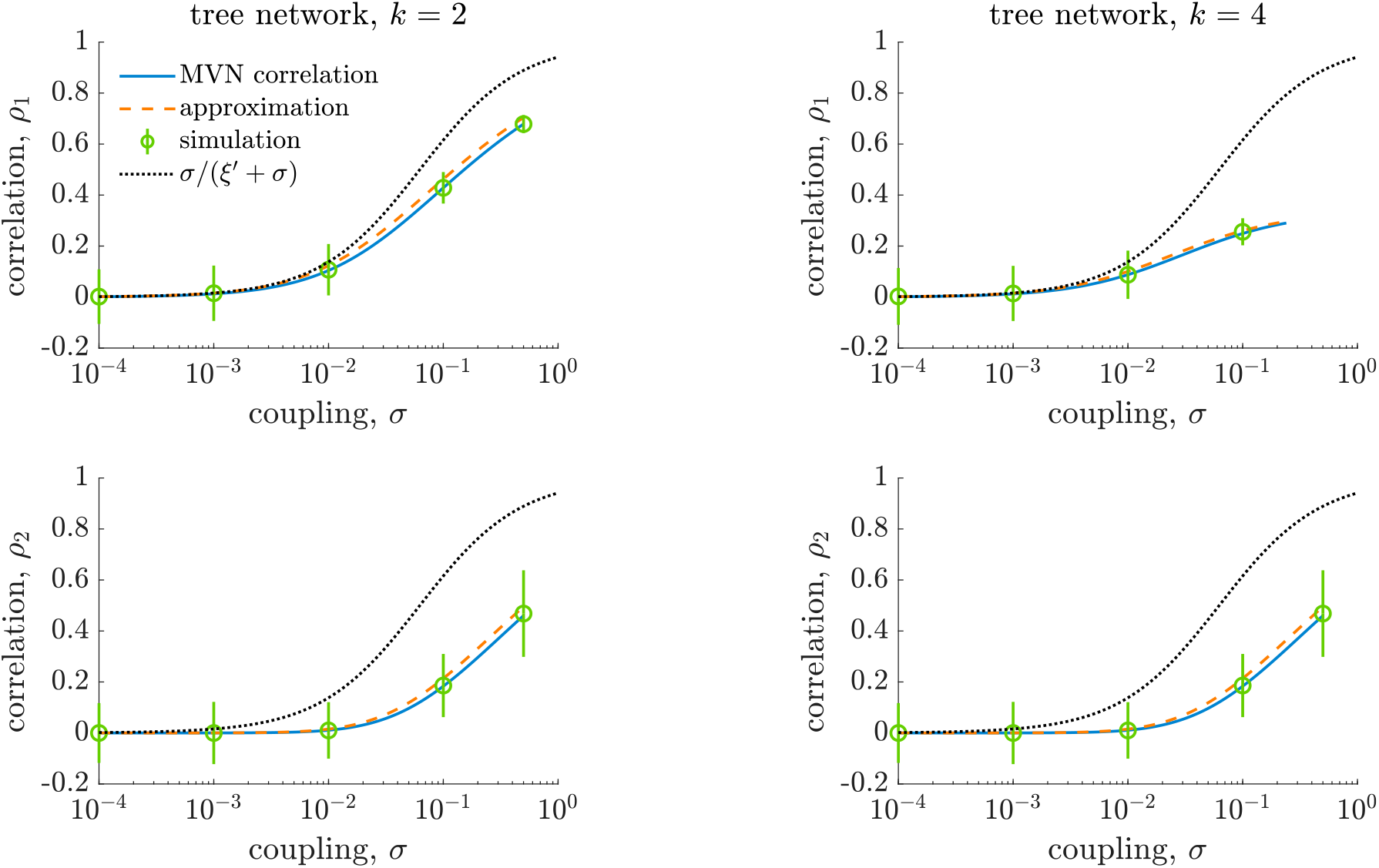
Comparing the MVN correlation *ρ_d_* and our approximation to stochastic simulations for a measles-like endemic disease in the UK in *T* populations arranged on the *D*-truncated *k*-regular tree network (*N* = 10^5^, *μ* = 5.5×10^−5^,*β* = 17/13, *ϵ* = 5.5×10^−5^,*γ* = 1/13). The coupling between interacting populations is *σ* ∈ [0,1/*k*]. The stochastic process is simulated on the *D*-truncated *k*-regular tree network, with *D* = 5 and *D* = 3 for *k* = 2,4, respectively. The process is simulated over a 200 year period using the Gillespie algorithm, with a burn-in period of 50 years, and generate 100 realisations of the process for each value of *σ*. The correlation is calculated as a time-weighted Pearson correlation coefficient for 50 ≤ *t* ≤ 200; error bars represent ±2 standard deviations.

### 3.3 The star network

#### 3.3.1 Network definition and notation

Finally, we consider the star network on *P* subpopulations, where there is a central ‘hub’ subpopulation (labelled as subpopulation 1) and *k* = *P* − 1 ‘leaf’ populations; there is no direct interaction between the leaf populations. A visualisation of the star network for *P* = 3 and *P* = 5 subpopulations is given in Figure 7. The coupling matrix **Σ** = (*σ_ij_*) is defined as

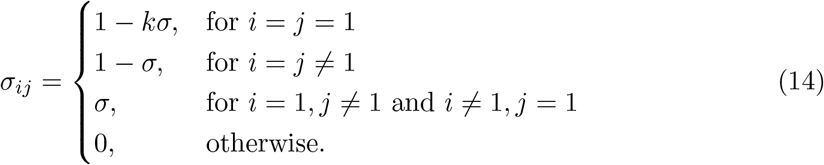

**Figure 7.**
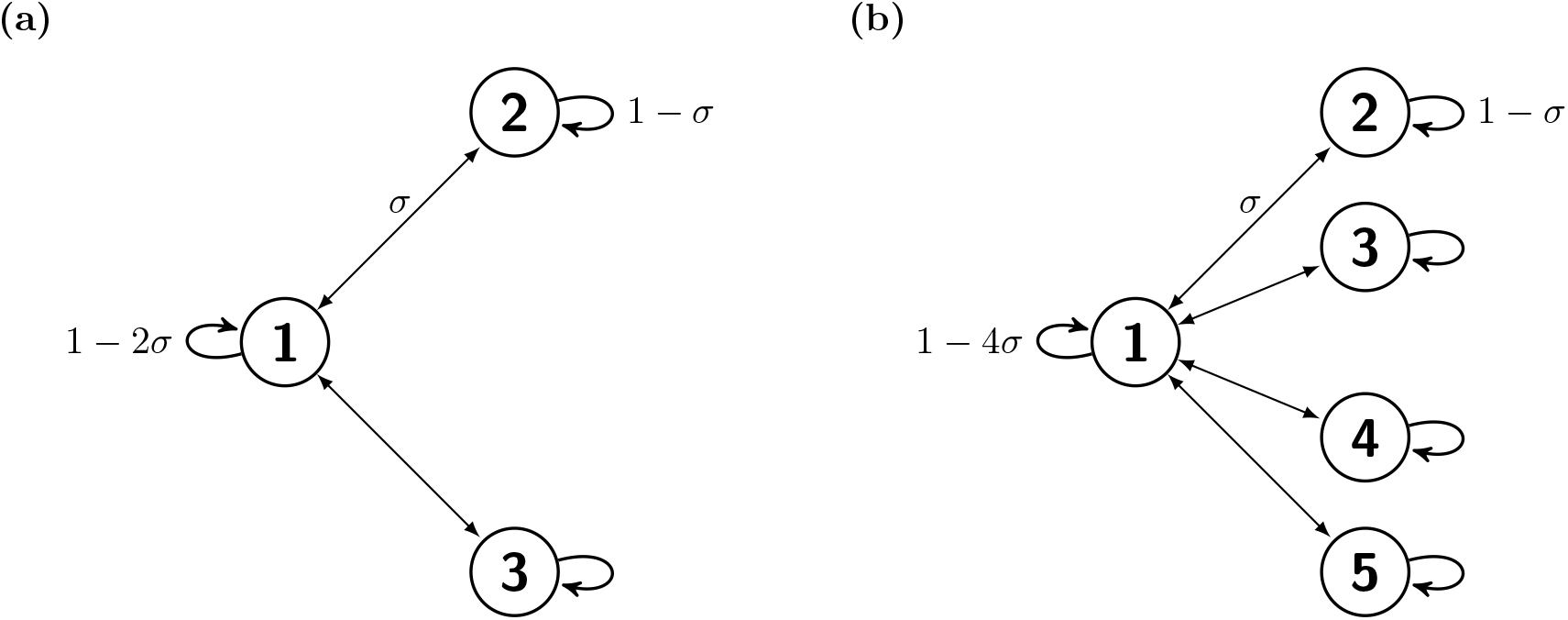
The star network on (a) *P* = 3 and (b) *P* = 5 populations. The coupling between any pair of neighbouring populations is *σ* ∈ [0,1/(*P* − 1)] and so the within-population coupling is 1 − (*P* − 1)*σ* for the hub population and 1 − *σ* for any leaf population.

Unlike the complete network and the *k*-regular tree network, the expected behaviour of the stochastic process is not the same within and between all subpopulations. This is because the hub subpopulation has *k* neighbours, whereas each leaf subpopulation has only one neighbour. However, we can still make some simplifications to the notation: the expected behaviour of the infection process is the same within any leaf subpopulation, or between any pair of leaf subpopulations, or between a leaf subpopulation and the hub subpopulation. We can therefore simplify our notation to distinguish between hub and leaf subpopulations. For the within-population moments, the superscript indicates whether the subpopulation is a hub (*H*) or a leaf (*L*) subpopulation:

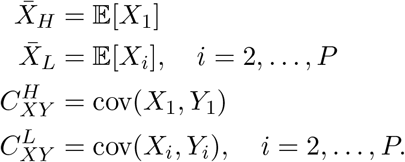

For the between-population moments, the superscript indicates whether one of the subpopulation is a hub (*H*) or if they are both leaf subpopulations (*L*); for 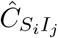 we distinguish between 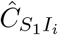 and 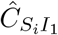:

Using the second-order moment closure approximation, the stochastic process on the star network for *P* subpopulations can be approximated by a set of seventeen ODEs: ten equations for the within-population moments, and seven equations for the between-population moments. These can be found in the Supplementary Information. We use these equations in both the analytic and the numerical results.

#### 3.3.2 Analytic results

For *P* identical subpopulations on the star network, we define the correlation between the number of infected individuals in the hub population and the number of infected individuals in a leaf population as

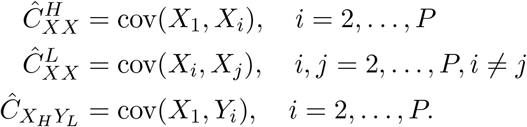

and the correlation between the number of infected individuals in two leaf subpopulations

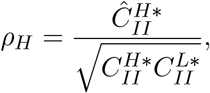

as

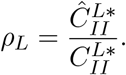

We can show that *ρ_H_* and *ρ_L_* are solution to the following pair of simultaneous equations:

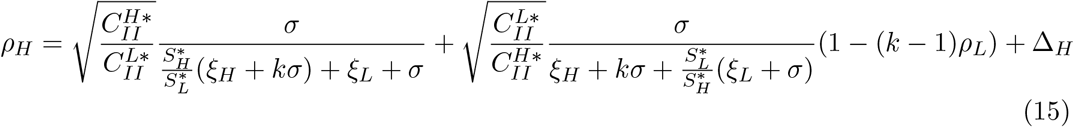

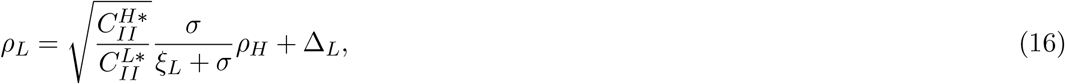

where

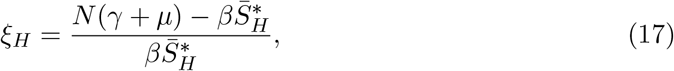

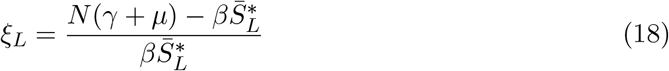

and

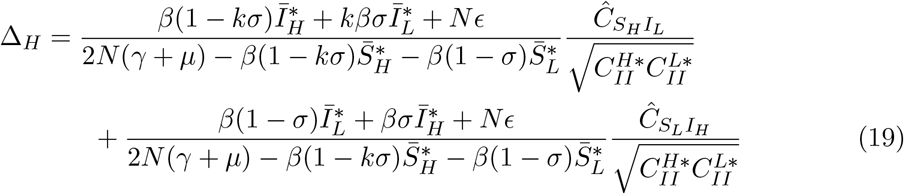

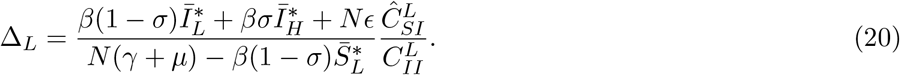

We derive this result by taking the moment equation for 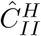 and 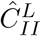 at equilibrium; full details of this derivation can be found in the Supplementary Information. Moreover, if Δ_*H*_, Δ_*L*_ ≪ 1 then we can further simplify this result to the following pair of simultaneous equations:

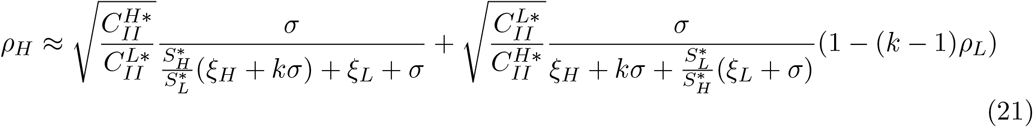

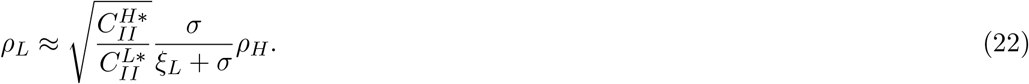

##### 3.3.3 Numerical results

We first numerically evaluate the effect of the number of leaf subpopulations *k* on the correlations *ρ_H_* and *ρ_L_* (Figure 8). Firstly, we note that, as with the complete and tree network, both *ρ_H_* and *ρ_L_* exhibit a sigmoidal shape, increasing from zero correlation from very low coupling. Secondly, the correlation between two leaf nodes is lower than between the hub and a leaf node; this is to be expected, as the leaf nodes are not directly connected to each other. Finally for a given coupling *σ* as the number of neighbours *k* increases then the correlation decreases; this holds for both *ρ_H_* and *ρ_L_*.

**Figure 8.**
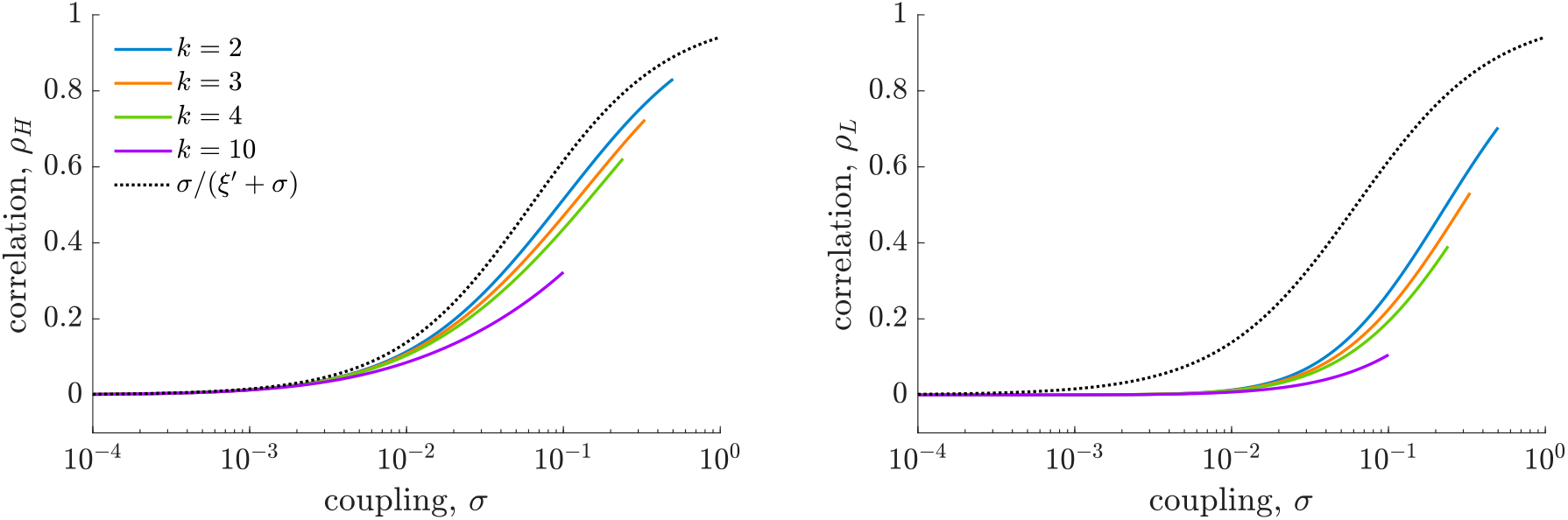
The effect of the number of leaf subpopulations *k* in the star network on the correlation between the number of infected individuals in the hub and a leaf population, *ρ_H_* (left), and two leaf populations, *ρ_L_* (right). Parameter values represent a measles-like endemic disease in the UK (*N* = 10^5^, *μ* = 5.5 × 10^−5^, *R*_0_ = 17, *ϵ* = 5.5 × 10^−5^, *γ* = 1/13). These values are calculated from the system of ODEs given in the Supplementary Information.

Next, we compare the MVN correlation and our approximation to the results of stochastic simulations (Figure 9). Firstly, we observe a close agreement between the MVN correlation and the stochastic simulations, which suggests that our use of the MVN moment closure approximation is justified. Secondly, there is little difference between the MVN correlation and our approximation (that is, Δ_*H*_ and Δ_*L*_ are small), and so our approximation is reasonable.

**Figure 9.**
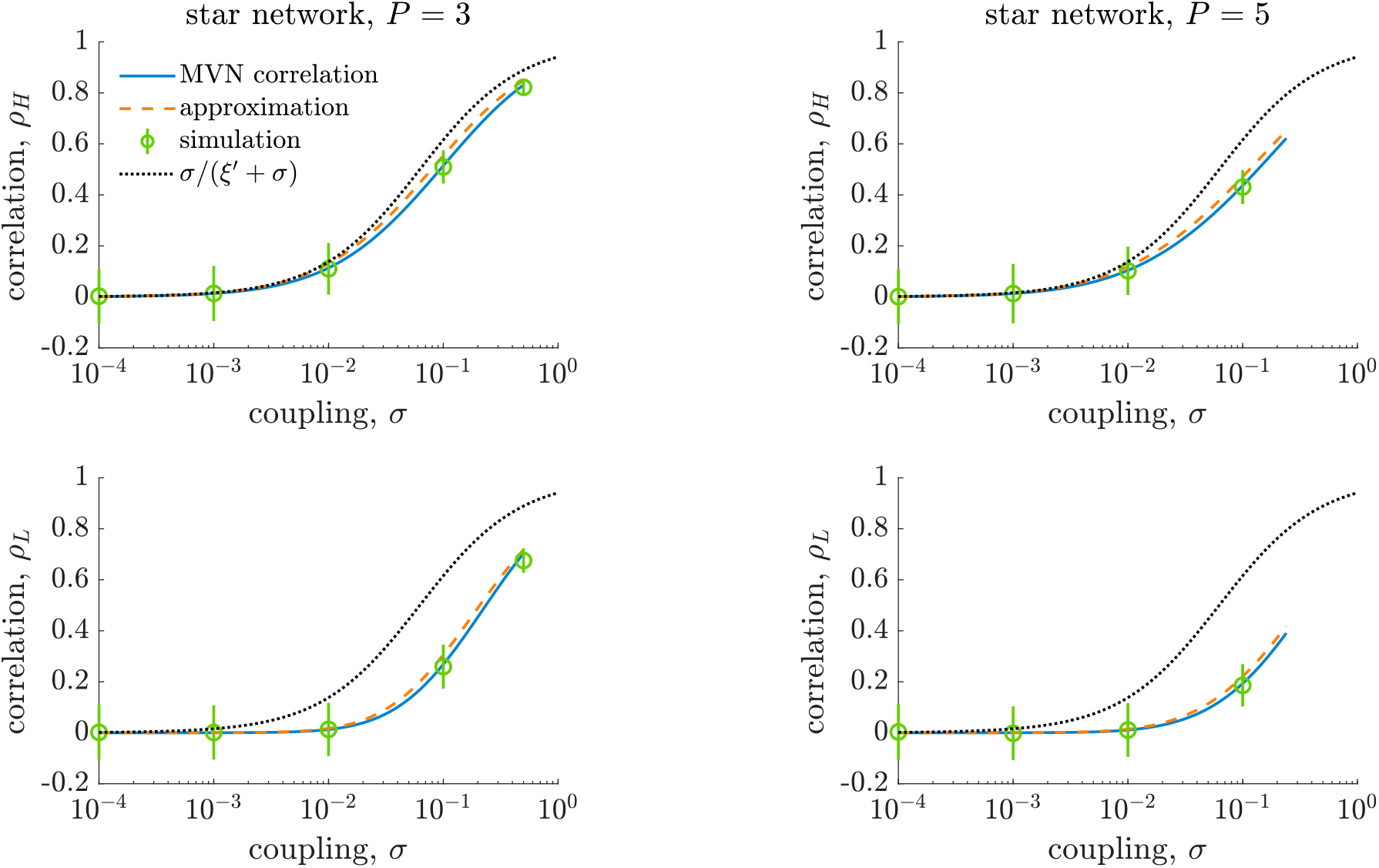
Comparing the analytic correlation, *ρ_H_* and *ρ_L_*, and our approximation to stochastic simulations for a measles-like endemic disease in the UK in *P* + 1 populations arranged on the star network (*N* = 10^5^, *μ* = 5.5 × 10^−5^, *β* = 17/13, *ϵ* = 5.5 × 10^−5^, *γ* = 1/13). The between-population coupling is fixed as *σ* ∈ [0,1] and within-population coupling is therefore 1 − *σ* in the hub population and 1 − *σ* in any leaf population. The stochastic process is simulated over a 200 year period using the Gillespie algorithm, with a burn-in period of 50 years, and generate 1000 realisations of the process for each value of *σ*. The correlation is calculated as a time-weighted Pearson correlation coefficient for 50 ≤ *t* ≤ 200; error bars represent ±2 standard deviations.

#### 3.4 Comparison of networks

We now compare our approximations to the correlation between the number of infected individuals in adjacent subpopulations for all three networks (Figure 10). All networks are chosen to have the same *k* external connections: the complete network with *P* = *k* + 1 populations, the *k*-regular tree network, and the star network with *P* = *k* + 1 populations. We observe that the correlation is highest in the complete network and lowest in the tree network. Moreover, the difference between the approximations increases as *k* increases.

**Figure 10.**
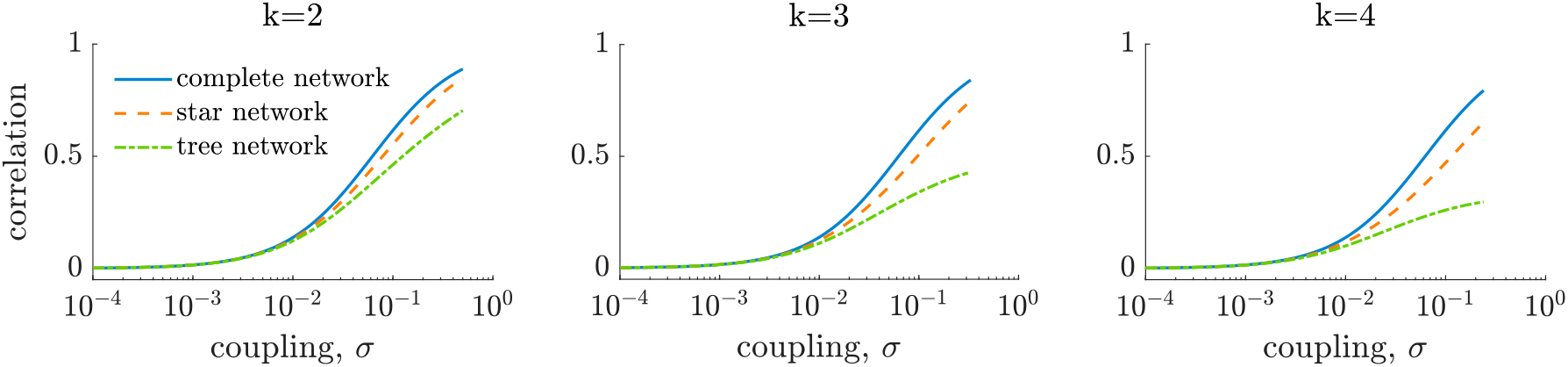
Comparison of our approximation to the correlation between a pair of adjacent populations in the complete network with *P* = *k* + 1 populations, the *k*-regular tree network and the star network with *P* = *k* + 1 populations.

We attribute this behaviour to the total number of neighbour subpopulations that the two focal subpopulations have, how many of those neighbours are common neighbours, and whether these common neighbours interact. As the total number of neighbours of each member of the focal pair increases then the correlation decreases; for a given total number of neighbours the correlation is higher when more of these neighbours are common between the two focal subpopulations, and is higher yet when these common neighbours also interact with each other.

For a given *k*, two focal subpopulations in the complete network and the star network both have a total of *k* – 1 subpopulations. In the star network, none of these subpopulations are common neighbours of the two focal subpopulations; however, in the complete network, all these subpopulations are common neighbours and all the common neighbours interact with each other, hence the correlation in the star network is lower than in the complete network. For the same *k*, two focal subpopulations in the *k*-regular tree network have twice the total number of neighbours compared to the star network and none of these neighbours are common neighbours for either network. As a result, the correlation is lower in the tree network than in the star network.

## 4 Discussion

A limitation of metapopulation models in epidemiological modelling is now to infer the coupling between subpopulations: existing models to not accurately describe human mobility in developing countries, such as Sub-Saharan Africa, and sufficiently detailed data on human mobility are often lacking. We propose that data on disease incidence can be used to infer the underlying coupling from observed correlations between subpopulations. We derive an approximation for the correlation *ρ* between the number of infected individuals in a given pair of subpopulations in certain network structures as a function of the coupling parameter *σ*. This provides a one-to-one mapping between the observable correlation *ρ* and the unknown coupling *σ*.

Our results extend the analysis of Meakin and Keeling (2018) from a simple two-population system to multiple populations arranged on a complete network, a *k*-regular tree network and a star network. Although we consider highly symmetric metapopulation networks, increased network complexity significantly reduces the analytic tractability of the model, compared to the two-population system. An alternative analytic relationship between the coupling and correlation has previously been derived for more general networks (Rozhnova et al., 2012); however, we believe that our results provide greater intuition and analytical traction.

In addition, these results improve our understanding of how metapopulation network structure affects endemic disease dynamics in the metapopulation as a whole, complementing existing research on epidemic diseases in metapopulation networks (Barthelemy et al., 2010; Lahodny and Allen, 2013; Wang and Wu, 2018; Yan et al., 2018). We find that network distance between subpopulations and network structure are key drivers of the correlation, although, surprisingly, in the complete network the correlation between any pair of subpopulations is independent of the total number of subpopulations. We hypothesise that the correlation between two given subpopulations is driven by the the number of neighbour subpopulations they both have, how many of these neighbours are shared between both subpopulations, and interactions between the neighbours.

Our research currently considers the mathematically tractable case of multiple identical populations on highly symmetric metapopulation networks. A natural extension of the our current results would be to allow heterogeneity in the transmission parameter β, or population size, although we have previously showed that heterogeneous population sizes significantly impact the tractability of the results (Meakin and Keeling, 2018). In addition, the simple network structures we consider here do not fully capture the observed characteristics of real-world spatial networks, such as heterogeneous population size, degree or edge weight (Guimerà et al., 2005; Colizza et al., 2006). We propose conducting a simulation-based study to examine in depth how the correlation between two focal subpopulations is affected by their neighbours, their neighbours’ neighbours and possible interactions between neighbours. This will allow us to elucidate which are the most important drivers of network correlations and overall endemic disease dynamics. A final limitation is that very few diseases are captured by the simple SIR compartmental model; however, it would be straightforward to extend the results presented here to more realistic models.

Our results provide a method by which the coupling can be estimated from the correlation between the number of infected individuals in two populations using data on disease incidence, allowing us to estimate the coupling between subpopulations even in the absence of mobility data. Our results also offer insight into the effect of metapopulation structure on endemic disease dynamics,

## 5 Conclusion

A limitation of metapopulation models in epidemiological modelling is now to infer the coupling between subpopulations. In this paper we relate the correlation between the number of infected individuals in two populations as a function of the coupling, considering systems of multiple identical interacting populations on highly-symmetric complex networks. Our results provide insight into the effect of metapopulation network structure on endemic disease dynamics and, used in combination with disease prevalence data, provide a method

## Author’s contributions

M.J.K. developed the initial concepts. S.R.M. performed the detailed mathematical analysis. Both authors played a role in writing and editing the manuscript.

