## Supplementary Information for "Correlations between stochastic endemic infection in multiple interacting subpopulations"

### Correlations between stochastic epidemics in multiple interacting subpopulations: supplementary information

#### 1 The ODE system approximating the stochastic epidemic metapopulation model on the complete network

For the stochastic epidemic metapopulation model on the complete network with $P$ populations, where the coupling between interacting populations is $\sigma \in [0, 1/k], k = P - 1$, we can approximate the stochastic process by the following system of 8 ODEs. This system comprises 5 equations for the within-population moments (of which two are first-order and three are second-order):

$$\frac{d\bar{S}}{dt} = \mu N - \frac{\beta}{N}(1-k\sigma)(C_{SI} + \bar{S}\bar{I}) - k\frac{\beta}{N}\sigma(\hat{C}_{SI} + \bar{S}\bar{I}) - \epsilon\bar{S} - \mu\bar{S} \tag{1}$$

$$\frac{d\bar{I}}{dt} = \frac{\beta}{N}(1-k\sigma)(C_{SI} + \bar{S}\bar{I}) + k\frac{\beta}{N}\sigma(\hat{C}_{SI} + \bar{S}\bar{I}) + \epsilon\bar{S} - \gamma\bar{I} - \mu\bar{I} \tag{2}$$

$$\frac{dC_{SS}}{dt} = \mu N\frac{\beta}{N}\bar{S}\bar{I} + \epsilon\bar{S} - \mu\bar{S} - 2\left(\frac{\beta}{N}\bar{I} + \epsilon + \mu\right) - \frac{\beta}{N}(1-k\sigma)(2\bar{S}-1) + k\frac{\beta}{N}\sigma(2\bar{S}-1)\hat{C}_{SI} \tag{3}$$

$$\begin{aligned}\frac{dC_{II}}{dt} &= \frac{\beta}{N}\bar{S}\bar{I} + \epsilon\bar{S} + (\gamma+\mu)\bar{I} + 2\left(\frac{\beta}{N}(1-k\sigma)\bar{S} - (\gamma+\mu)\right)C_{II} \\ &\quad + \left(\frac{\beta}{N}(1-k\sigma)(2\bar{I}+1) + 2k\frac{\beta}{N}\sigma\bar{I} + 2\epsilon\right)C_{SI} + 2k\frac{\beta}{N}\sigma\bar{S}\hat{C}_{II} + k\frac{\beta}{N}\sigma\hat{C}_{SI}\end{aligned} \tag{4}$$

$$\begin{aligned}\frac{dC_{SI}}{dt} &= -\frac{\beta}{N}\bar{S}\bar{I} - \epsilon\bar{S} - \mu\bar{I} + \left(\frac{\beta}{N}\bar{I} + \epsilon\right)C_{SS} - \frac{\beta}{N}(1-k\sigma)\bar{S}C_{II} \\ &\quad + \left(\frac{\beta}{N}(1-k\sigma)(\bar{S}-\bar{I}-1) - k\frac{\beta}{N}\sigma\bar{I} - \epsilon - \gamma - 2\mu\right)C_{SI} - k\frac{\beta}{N}\sigma\bar{S}\hat{C}_{II} + k\frac{\beta}{N}\sigma(\bar{S}-1)\hat{C}_{SI},\end{aligned} \tag{5}$$

(6)

and 3 equations for the between-population moments:

$$\frac{d\hat{C}_{SS}}{dt} = -2\frac{\beta}{N}\sigma\bar{S}C_{SI} - 2\left(\frac{\beta}{N}\bar{I} + \epsilon + \mu\right)\hat{C}_{SS} - 2\left(\frac{\beta}{N}(1-k\sigma)\bar{S} + (k-1)\frac{\beta}{N}\sigma\bar{S}\right)\hat{C}_{SI} \tag{7}$$

$$\frac{d\hat{C}_{II}}{dt} = 2\frac{\beta}{N}\sigma\bar{S}C_{II} + 2\left(\frac{\beta}{N}(1-k\sigma)\bar{S} + (k-1)\frac{\beta}{N}\sigma\bar{S} - \gamma - \mu\right)\hat{C}_{II} + 2\left(\frac{\beta}{N}\bar{I} + \epsilon\right)\hat{C}_{SI} \tag{8}$$

$$\begin{aligned}\frac{d\hat{C}_{SI}}{dt} &= -\frac{\beta}{N}\sigma\bar{S}C_{II} + \frac{\beta}{N}\sigma\bar{S}C_{SI} + \left(\frac{\beta}{N}\bar{I} + \epsilon\right)\hat{C}_{SS} - \left(\frac{\beta}{N}(1-k\sigma) + (k-1)\frac{\beta}{N}\sigma\bar{S}\right)\hat{C}_{II} \\ &\quad + \left(\frac{\beta}{N}(1-k\sigma)(\bar{S}-\bar{I}) + \frac{\beta}{N}\sigma((k-1)\bar{S} - k\bar{I}) - \epsilon - \gamma - 2\mu\right)\hat{C}_{SI}.\end{aligned} \tag{9}$$

#### 2 Derivation of the approximation for the complete network

For the complete network on $P$ populations, where the coupling between interacting populations is $\sigma \in [0, 1/k], k = P - 1$, we can show that the correlation, $\rho$, between the number of infected individuals in any pair of populations is equal to

$$\rho = \frac{\sigma}{\xi + \sigma} - \Delta, \tag{10}$$

where

$$\xi = \frac{N(\gamma + \mu) - \beta \bar{S}^*}{\beta \bar{S}^*} \tag{11}$$

and

$$\Delta = \frac{(\beta \bar{I}^* + N\epsilon)\frac{\hat{C}^*_{SI}}{C^*_{II}}}{\beta(1-\sigma)\bar{S}^* - N(\gamma + \mu)}. \tag{12}$$

To derive this result we begin with the moment equation for $\hat{C}_{II}$:

$$\frac{d\hat{C}_{II}}{dt} = 2\frac{\beta}{N}\sigma \bar{S} C_{II} + 2\left(\frac{\beta}{N}(1-\sigma)\bar{S} - \gamma - \mu\right)\hat{C}_{II} + 2\left(\frac{\beta}{N}\bar{I} + \epsilon\right)\hat{C}_{SI}. \tag{13}$$

At equilibrium, $d\hat{C}_{II}/dt = 0$ and if we divide by $2C^*_{II}/N$, then

$$0 = \beta\sigma\bar{S}^* + \big(\beta(1-\sigma)\bar{S}^* - N(\gamma+\mu)\big)\,\rho + (\beta\bar{I}^* + N\epsilon)\frac{\hat{C}^*_{SI}}{C^*_{II}}, \tag{14}$$

and hence we have the following approximation for the correlation:

$$\begin{aligned}
\rho &= \frac{-\beta\sigma\bar{S}^*}{\beta(1-\sigma)\bar{S}^* - N(\gamma+\mu)} - \frac{\beta\bar{I}^* + N\epsilon}{\beta(1-\sigma)\bar{S}^* - N(\gamma+\mu)}\frac{\hat{C}^*_{SI}}{C^*_{II}} \\
&= \frac{\sigma}{\frac{N(\gamma+\mu)-\beta\bar{S}^*}{\beta\bar{S}^*} + \sigma} - \Delta \\
&= \frac{\sigma}{\xi + \sigma} - \Delta.
\end{aligned} \tag{15}$$

#### 3 The approximation for the MVN correlation on the complete network is independent of the number of populations $P$

For the stochastic epidemic metapopulation model on the complete network with $P = k + 1$ populations where the coupling between any pair of populations is $\sigma \in [0, 1/k]$, then the MVN correlation can be approximated by

$$\rho = \frac{\sigma}{\xi' + \sigma}, \tag{16}$$

where $\xi' = \epsilon/(\mu(R_0 - 1))$. This result is independent of the number of populations $P$.

##### 3.1 Network definition and notation

We consider a metapopulation network on three populations with coupling matrix $\mathbf{\Sigma} = (\sigma_{ij})_{ij}$ defined as

$$\sigma_{ij} = \begin{cases} 1 - \sigma - a, & \text{for } i = j = 1, 2 \\ 1 - 2a, & \text{for } i = j = 3 \\ \sigma, & \text{for } (i, j) \in \{(1, 2), (2, 1)\} \\ a, & \text{for } (i, j) \in \{(1, 3), (3, 1), (2, 3), (3, 2)\}, \end{cases} \tag{17}$$

where $\sigma \in [0, 1]$ and $a \in [0, \sigma]$. A visual representation of this metapopulation is given in Figure S1.

In this metapopulation all subpopulations are epidemiologically identical (that is, they are the same size and have identical epidemiological parameters). In addition, subpopulations 1 and 2 are isomorphic in the weighted network, but are only isomorphic with subpopulation 3 when $a = \sigma$. We retain our original notation, using $\bar{X}_i = \mathbb{E}[X_i]$, $C_{X_iY_i} = \text{Cov}(X_i, Y_i)$ and $\hat{C}_{X_iY_j} = \text{Cov}(X_iY_j)$.

##### 3.2 Approximation for correlation between subpopulation 1 and subpopulation 2

The correlation between subpopulation 1 and subpopulation 2 is given by

$$\rho_{12} = \frac{\hat{C}_{I_1I_2}}{\sqrt{C_{I_1I_1}C_{I_2I_2}}} = \frac{\hat{C}_{I_1I_2}}{C_{I_2I_2}}. \tag{18}$$

To derive an approximation for this, we begin with the moment equation for $\hat{C}_{I_1I_2}$:

$$\frac{d\hat{C}_{I_1I_2}}{dt} = 2\Bigg[\left(\frac{\beta}{N}(1 - \sigma - a)\bar{S}_1 - (\gamma + \mu)\right)\hat{C}_{I_1I_2} + \frac{\beta}{N}\sigma\bar{S}_1C_{I_2I_2} + \frac{\beta}{N}a\bar{S}_1\hat{C}_{I_2I_3} + \left(\frac{\beta}{N}(1 - \sigma - a)\bar{I}_1 + \frac{\beta}{N}\sigma\bar{I}_2 + \frac{\beta}{N}a\bar{I}_3 + \epsilon\right)\hat{C}_{S_1I_2}\Bigg]. \tag{19}$$

At equilibrium $d\hat{C}_{I_1I_2}/dt = 0$, and if we divide by $2C_{I_2I_2}/N$ then

$$0 = [\beta(1 - \sigma - a)\bar{S}_1 - N(\gamma + \mu)]\rho_{12} + \beta\sigma\bar{S}_1 + \beta a\bar{S}_1\sqrt{\frac{V_3}{V_1}}\rho_{23} + \big(\beta(1 - \sigma - a)\bar{I}_1 + \beta\sigma\bar{I}_2 + \beta a\bar{I}_3 + \epsilon\big)\frac{\hat{C}_{S_1I_2}}{C_{I_2I_2}} \tag{20}$$

$$= [\beta(1 - \sigma)\bar{S}_1 - N(\gamma + \mu)]\rho_{12} + \beta\sigma\bar{S}_1 + \beta a\bar{S}_1\left(\sqrt{\frac{V_3}{V_1}}\rho_{23} - \rho_{12}\right) + \big(\beta(1 - \sigma - a)\bar{I}_1 + \beta\sigma\bar{I}_2 + \beta a\bar{I}_3 + \epsilon\big)\frac{\hat{C}_{S_1I_2}}{C_{I_2I_2}}, \tag{21}$$

and hence we have the following approximation for the correlation between subpopulation 1 and subpopulation 2:

$$\rho_{12} = \frac{\beta\sigma\bar{S}_1}{N(\gamma+\mu)-\beta(1-\sigma)\bar{S}_1} + \frac{\beta a\bar{S}_1}{N(\gamma+\mu)-\beta(1-\sigma)\bar{S}_1}\left(\sqrt{\frac{V_3}{V_1}}\rho_{23}-\rho_{12}\right) - \Delta \tag{22}$$

$$= \frac{\sigma}{\xi+\sigma} + \frac{a}{\xi+\sigma}\left(\sqrt{\frac{V_3}{V_1}}\rho_{23}-\rho_{12}\right) - \Delta, \tag{23}$$

where $\Delta$ is a correction term:

$$\Delta = \frac{(\beta(1-\sigma-a)\bar{I}_1 + \beta\sigma\bar{I}_2 + \beta a\bar{I}_3 + \epsilon)}{\beta(1-\sigma)\bar{S}_1 - N(\gamma+\mu)}\frac{\hat{C}_{S_1I_2}}{C_{I_2I_2}}. \tag{24}$$

If $\Delta \ll 1$, then we have the following simplified expression for the correlation:

$$\rho_{12} \approx \frac{\sigma}{\xi+\sigma} + \frac{a}{\xi+\sigma}\left(\sqrt{\frac{V_3}{V_1}}\rho_{23}-\rho_{12}\right). \tag{25}$$

We consider how the second term of Equation 25 changes for $a \in [0,\sigma]$. When $a = 0$ then the second term vanishes and $\rho_{12} = \sigma/(\xi'+\sigma)$. When $a = \sigma$ then $V_1 = V_3$ and $\rho_{12} = \rho_{23}$, so $(\sqrt{V_3/V_1}\rho_{23} - \rho_{12}) = 0$ and thus we also have $\rho_{12} = \sigma/(\xi'+\sigma)$.

When $a \in (0,\sigma)$ then $0 < \rho_{23} < \rho_{12}$ and so $(\sqrt{V_3/V_1}\rho_{23} - \rho_{12}) < 0$ if and only if $\sqrt{V_3/V_1} < \rho_{12}/\rho_{23}$.

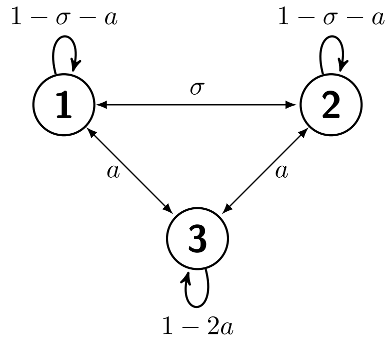


**Figure S1.** A metapopulation on three identical subpopulations. The coupling between populations 1 and 2 is $\sigma \in [0,1]$, and the coupling between populations 1 and 3, or 2 and 3, is $a \in [0,\sigma]$.

### 4 Discussions of assumptions of dynamics on the $D$-truncated $k$-regular tree network

When defining the $D$-truncated $k$-regular tree network and deriving the set of ODEs that approximate the stochastic epidemic process on the network, we make three assumptions. To justify these assumptions, we detail the meaning and result of each assumption, and the conditions under which the assumption is likely to hold.

**Assumption 1: covariances are negligibly small for subpopulations that are far apart.** We assume that for $D$ sufficiently large, $\hat{C}_{XY}^{(d)} = 0, \forall d > D$. The result of this is that we can write down a smaller and simpler set of ODEs that approximate the stochastic process on the $D$-truncated $k$-regular tree network: without this assumption, we would also have to write down ODEs for covariances $\hat{C}_{XY}^{(d)}$ where $D < d \leq 2D$.

**Assumption 2: covariances between any pair of subpopulations the same distance apart are the same.** We maintain the simplification from the full $k$-regular tree network that $\hat{C}_{XY}^{(d)}$ is the same for any pair of subpopulations distance $d$ apart. Although this is not true (the covariances between adjacent populations at the centre of the truncated tree network will be different to covariances between adjacent subpopulations at the edge of the network, for example), in practice it will have little effect on the final results: SOMETHING. This assumption also reduces the total number of ODEs we need to write down to approximate the stochastic process.

**Assumption 3: the behaviour of the moments in the truncated tree network is the same as in the full tree network.** We assume that the expected behaviour of the first- and second-order central moments in the origin node ($\bar{X}$ and $C_{XY}$) in the $D$-truncated $k$-regular tree network will be the same as in the full $k$-regular tree network; we also assume that the expected behaviour of the second-order moments between the origin node and subpopulations distance $d \ll D$ ($\hat{C}_{XY}$) will be the same as in the full $k$-regular tree network.

### 5 The ODE system approximating the stochastic epidemic metapopulation model on the tree network

#### 5.1 The full $k$-regular tree network

We can approximate the stochastic epidemic process on the full $k$-regular tree network, where the coupling between interacting populations is $\sigma \in [0, 1/k]$, by the following system of ODES. This system comprises 5 equations for the within-population moments (of which two are first-order and three are second-order):

$$\frac{d\bar{S}}{dt} = \mu N - \frac{\beta}{N}(1-k\sigma)(C_{SI} + \bar{S}\bar{I}) - k\frac{\beta}{N}\sigma(\hat{C}^{(1)}_{SI} + \bar{S}\bar{I}) - \epsilon\bar{S} - \mu\bar{S} \tag{26}$$

$$\frac{d\bar{I}}{dt} = \frac{\beta}{N}(1-k\sigma)(C_{SI} + \bar{S}\bar{I}) + k\frac{\beta}{N}\sigma(\hat{C}^{(1)}_{SI} + \bar{S}\bar{I}) + \epsilon\bar{S} - \gamma\bar{I} - \mu\bar{I} \tag{27}$$

$$\frac{dC_{SS}}{dt} = \mu N\frac{\beta}{N}\bar{S}\bar{I} + \epsilon\bar{S} - \mu\bar{S} - 2\left(\frac{\beta}{N}\bar{I} + \epsilon + \mu\right) - \frac{\beta}{N}(1-k\sigma)(2\bar{S}-1) + k\frac{\beta}{N}\sigma(2\bar{S}-1)\hat{C}^{(1)}_{SI} \tag{28}$$

$$\begin{aligned}\frac{dC_{II}}{dt} &= \frac{\beta}{N}\bar{S}\bar{I} + \epsilon\bar{S} + (\gamma+\mu)\bar{I} + 2\left(\frac{\beta}{N}(1-k\sigma)\bar{S} - (\gamma+\mu)\right)C_{II} \\ &\quad + \left(\frac{\beta}{N}(1-k\sigma)(2\bar{I}+1) + 2k\frac{\beta}{N}\sigma\bar{I} + 2\epsilon\right)C_{SI} + 2k\frac{\beta}{N}\sigma\bar{S}\hat{C}^{(1)}_{II} + k\frac{\beta}{N}\sigma\hat{C}^{(1)}_{SI}\end{aligned} \tag{29}$$

$$\begin{aligned}\frac{dC_{SI}}{dt} &= -\frac{\beta}{N}\bar{S}\bar{I} - \epsilon\bar{S} - \mu\bar{I} + \left(\frac{\beta}{N}\bar{I} + \epsilon\right)C_{SS} - \frac{\beta}{N}(1-k\sigma)\bar{S}C_{II} \\ &\quad + \left(\frac{\beta}{N}(1-k\sigma)(\bar{S}-\bar{I}-1) - k\frac{\beta}{N}\sigma\bar{I} - \epsilon - \gamma - 2\mu\right)C_{SI} - k\frac{\beta}{N}\sigma\bar{S}\hat{C}^{(1)}_{II} + k\frac{\beta}{N}\sigma(\bar{S}-1)\hat{C}^{(1)}_{SI},\end{aligned} \tag{30}$$

and $3d$ equations for the between-population moments, $d \geq 1$:

$$\frac{d\hat{C}^{(d)}_{SS}}{dt} = -2\frac{\beta}{N}\sigma\bar{S}\hat{C}^{(d-1)}_{SI} - 2\left(\frac{\beta}{N}\bar{I} + \epsilon + \mu\right)\hat{C}^{(d)}_{SS} - 2\frac{\beta}{N}(1-k\sigma)\bar{S}\hat{C}^{(d)}_{SI} - 2(k-1)\frac{\beta}{N}\sigma\bar{S}\hat{C}^{(d+1)}_{SI} \tag{31}$$

$$\frac{d\hat{C}^{(d)}_{II}}{dt} = 2\frac{\beta}{N}\sigma\bar{S}\hat{C}^{(d-1)}_{II} + 2\left(\frac{\beta}{N}(1-k\sigma)\bar{S} - \gamma - \mu\right)\hat{C}^{(d)}_{II} + 2\left(\frac{\beta}{N}\bar{I} + \epsilon\right)\hat{C}^{(d)}_{SI} + 2(k-1)\frac{\beta}{N}\sigma\bar{S}\hat{C}^{(d+1)}_{II} \tag{32}$$

$$\begin{aligned}\frac{d\hat{C}^{(d)}_{SI}}{dt} &= -\frac{\beta}{N}\sigma\bar{S}C^{(d-1)}_{II} + \frac{\beta}{N}\sigma\bar{S}C^{(d-1)}_{SI} + \left(\frac{\beta}{N}\bar{I} + \epsilon\right)\hat{C}^{(d)}_{SS} - \frac{\beta}{N}(1-k\sigma)\hat{C}^{(d)}_{II} \\ &\quad + \left(\frac{\beta}{N}(1-k\sigma)(\bar{S}-\bar{I}) - k\frac{\beta}{N}\sigma\bar{I} - \epsilon - \gamma - 2\mu\right)\hat{C}^{(d)}_{SI} - (k-1)\frac{\beta}{N}\sigma\bar{S}\hat{C}^{(d+1)}_{II} + (k-1)\frac{\beta}{N}\sigma\bar{S}\hat{C}^{(d+1)}_{SI},\end{aligned} \tag{33}$$

and where $\hat{C}^{(0)}_{XY} = C_{XY}$.

#### 5.2 The $D$-truncated $k$-regular tree network

We can approximate the stochastic epidemic process on the $D$-truncated $k$-regular tree network, where the coupling between interacting populations is $\sigma \in [0, 1/k]$, by the following system of $3D + 5$ ODEs. This system comprises

5 equations for the within-population moments (of which two are first-order and three are second-order):

$$\frac{d\bar{S}}{dt} = \mu N - \frac{\beta}{N}(1-k\sigma)(C_{SI}+\bar{S}\bar{I}) - k\frac{\beta}{N}\sigma(\hat{C}_{SI}^{(1)}+\bar{S}\bar{I}) - \epsilon\bar{S} - \mu\bar{S} \tag{34}$$

$$\frac{d\bar{I}}{dt} = \frac{\beta}{N}(1-k\sigma)(C_{SI}+\bar{S}\bar{I}) + k\frac{\beta}{N}\sigma(\hat{C}_{SI}^{(1)}+\bar{S}\bar{I}) + \epsilon\bar{S} - \gamma\bar{I} - \mu\bar{I} \tag{35}$$

$$\frac{dC_{SS}}{dt} = \mu N\frac{\beta}{N}\bar{S}\bar{I} + \epsilon\bar{S} - \mu\bar{S} - 2\left(\frac{\beta}{N}\bar{I}+\epsilon+\mu\right) - \frac{\beta}{N}(1-k\sigma)(2\bar{S}-1) + k\frac{\beta}{N}\sigma(2\bar{S}-1)\hat{C}_{SI}^{(1)} \tag{36}$$

$$\begin{aligned}\frac{dC_{II}}{dt} &= \frac{\beta}{N}\bar{S}\bar{I} + \epsilon\bar{S} + (\gamma+\mu)\bar{I} + 2\left(\frac{\beta}{N}(1-k\sigma)\bar{S} - (\gamma+\mu)\right)C_{II} \\ &\quad + \left(\frac{\beta}{N}(1-k\sigma)(2\bar{I}+1) + 2k\frac{\beta}{N}\sigma\bar{I} + 2\epsilon\right)C_{SI} + 2k\frac{\beta}{N}\sigma\bar{S}\hat{C}_{II}^{(1)} + k\frac{\beta}{N}\sigma\hat{C}_{SI}^{(1)}\end{aligned} \tag{37}$$

$$\begin{aligned}\frac{dC_{SI}}{dt} &= -\frac{\beta}{N}\bar{S}\bar{I} - \epsilon\bar{S} - \mu\bar{I} + \left(\frac{\beta}{N}\bar{I}+\epsilon\right)C_{SS} - \frac{\beta}{N}(1-k\sigma)\bar{S}C_{II} \\ &\quad + \left(\frac{\beta}{N}(1-k\sigma)(\bar{S}-\bar{I}-1) - k\frac{\beta}{N}\sigma\bar{I} - \epsilon - \gamma - 2\mu\right)C_{SI} - k\frac{\beta}{N}\sigma\bar{S}\hat{C}_{II}^{(1)} + k\frac{\beta}{N}\sigma(\bar{S}-1)\hat{C}_{SI}^{(1)},\end{aligned} \tag{38}$$

and $3D$ equations for the between-population moments; for $d = 1, \ldots, D-1$ we have:

$$\frac{d\hat{C}_{SS}^{(d)}}{dt} = -2\frac{\beta}{N}\sigma\bar{S}\hat{C}_{SI}^{(d-1)} - 2\left(\frac{\beta}{N}\bar{I}+\epsilon+\mu\right)\hat{C}_{SS}^{(d)} - 2\frac{\beta}{N}(1-k\sigma)\bar{S}\hat{C}_{SI}^{(d)} - 2(k-1)\frac{\beta}{N}\sigma\bar{S}\hat{C}_{SI}^{(d+1)} \tag{39}$$

$$\frac{d\hat{C}_{II}^{(d)}}{dt} = 2\frac{\beta}{N}\sigma\bar{S}\hat{C}_{II}^{(d-1)} + 2\left(\frac{\beta}{N}(1-k\sigma)\bar{S}-\gamma-\mu\right)\hat{C}_{II}^{(d)} + 2\left(\frac{\beta}{N}\bar{I}+\epsilon\right)\hat{C}_{SI}^{(d)} + 2(k-1)\frac{\beta}{N}\sigma\bar{S}\hat{C}_{II}^{(d+1)} \tag{40}$$

$$\begin{aligned}\frac{d\hat{C}_{SI}^{(d)}}{dt} &= -\frac{\beta}{N}\sigma\bar{S}C_{II}^{(d-1)} + \frac{\beta}{N}\sigma\bar{S}C_{SI}^{(d-1)} + \left(\frac{\beta}{N}\bar{I}+\epsilon\right)\hat{C}_{SS}^{(d)} - \frac{\beta}{N}(1-k\sigma)\hat{C}_{II}^{(d)} \\ &\quad + \left(\frac{\beta}{N}(1-k\sigma)(\bar{S}-\bar{I}) - k\frac{\beta}{N}\sigma\bar{I} - \epsilon - \gamma - 2\mu\right)\hat{C}_{SI}^{(d)} - (k-1)\frac{\beta}{N}\sigma\bar{S}\hat{C}_{II}^{(d+1)} + (k-1)\frac{\beta}{N}\sigma\bar{S}\hat{C}_{SI}^{(d+1)},\end{aligned} \tag{41}$$

and for $d = D$ we have:

$$\frac{d\hat{C}_{SS}^{(D)}}{dt} = -2\frac{\beta}{N}\sigma\bar{S}\hat{C}_{SI}^{(D-1)} - 2\left(\frac{\beta}{N}\bar{I}+\epsilon+\mu\right)\hat{C}_{SS}^{(D)} - 2\frac{\beta}{N}(1-k\sigma)\bar{S}\hat{C}_{SI}^{(D)} \tag{42}$$

$$\frac{d\hat{C}_{II}^{(D)}}{dt} = 2\frac{\beta}{N}\sigma\bar{S}\hat{C}_{II}^{(D-1)} + 2\left(\frac{\beta}{N}(1-k\sigma)\bar{S}-\gamma-\mu\right)\hat{C}_{II}^{(D)} + 2\left(\frac{\beta}{N}\bar{I}+\epsilon\right)\hat{C}_{SI}^{(D)} \tag{43}$$

$$\begin{aligned}\frac{d\hat{C}_{SI}^{(D)}}{dt} &= -\frac{\beta}{N}\sigma\bar{S}C_{II}^{(D-1)} + \frac{\beta}{N}\sigma\bar{S}C_{SI}^{(D-1)} + \left(\frac{\beta}{N}\bar{I}+\epsilon\right)\hat{C}_{SS}^{(D)} - \frac{\beta}{N}(1-k\sigma)\hat{C}_{II}^{(D)} \\ &\quad + \left(\frac{\beta}{N}(1-k\sigma)(\bar{S}-\bar{I}) - k\frac{\beta}{N}\sigma\bar{I} - \epsilon - \gamma - 2\mu\right)\hat{C}_{SI}^{(D)}.\end{aligned} \tag{44}$$

#### 6 Derivation of the approximation for the $k$-regular tree network

For the $k$-regular tree network, where the coupling between interacting populations is $\sigma \in [0, 1/k], k = P - 1$, we can show that the correlation, $\rho$, between the number of infected individuals in any pair of populations distance $d$ apart is the solution to

$$\rho_d = \frac{\sigma}{\xi + k\sigma} \left(\rho_{d-1} + (k-1)\rho_{d+1}\right) - \Delta^{(d)}, \tag{45}$$

where

$$\xi = \frac{N(\gamma + \mu) - \beta \bar{S}^*}{\beta \bar{S}^*} \tag{46}$$

and

$$\Delta^{(d)} = \frac{(\beta \bar{I}^* + N\epsilon)}{\beta(1 - k\sigma)\bar{S}^* - N(\gamma + \mu)} \frac{\hat{C}_{SI}^{(d)*}}{C_{II}^*}. \tag{47}$$

We derive this result we begin with the moment equation $\hat{C}_{II}^{(d)}$:

$$\begin{aligned}\frac{\hat{C}_{II}^{(d)}}{dt} &= 2\frac{\beta}{N}\sigma \bar{S}\hat{C}_{II}^{(d-1)} + 2\left(\frac{\beta}{N}(1 - k\sigma)\bar{S} - (\gamma + \mu)\right)\hat{C}_{II}^{(d)} + 2\left(\frac{\beta}{N}\bar{I} + \epsilon\right)\hat{C}_{SI}^{(d)} \\ &\quad + 2(k-1)\frac{\beta}{N}\sigma \bar{S}\hat{C}_{II}^{(d+1)}\end{aligned} \tag{48}$$

At equilibrium $d\hat{C}_{II}^{(d)}/dt = 0$ and if we divide by $2C_{II}^*/N$ then

$$\begin{aligned}0 &= \beta\sigma\bar{S}^*\rho_{d-1} + (\beta(1 - k\sigma)\bar{S}^* - N(\gamma + \mu))\rho_d + \beta\sigma\bar{S}^*(k-1)\rho_{d+1} \\ &\quad + (\beta\bar{I}^* + N\epsilon)\frac{\hat{C}_{SI}^{(d)*}}{C_{II}^*},\end{aligned} \tag{49}$$

and hence we have the following relationship between $\rho_{d-1}, \rho_d$ and $\rho_{d+1}$

$$\begin{aligned}\rho_d &= \frac{\beta\sigma\bar{S}^*}{N(\gamma + \mu) - \beta(1 - k\sigma)\bar{S}^*}\left(\rho_{d-1} + (k-1)\rho_{d+1}\right) - \frac{\beta\bar{I}^* + N\epsilon}{N(\gamma + \mu) - \beta(1 - k\sigma)\bar{S}^*}\frac{\hat{C}_{SI}^{(d)*}}{C_{II}^*} \\ &= \frac{\sigma}{\frac{N(\gamma+\mu) - \beta\bar{S}^*}{\beta\bar{S}^*} + k\sigma}\left(\rho_{d-1} + (k-1)\rho_{d+1}\right) - \Delta_k^{(d)} \\ &= \frac{\sigma}{\xi + k\sigma}\left(\rho_{d-1} + (k-1)\rho_{d+1}\right) - \Delta^{(d)}.\end{aligned} \tag{50}$$

#### 7 The ODE system approximating the stochastic epidemic metapopulation model on the star network

For the stochastic epidemic process on the star network with $k$ leaf populations, where the coupling between interacting populations is $\sigma \in [0, 1/k]$, by the following system of seventeen ODES. This system comprises ten equations for the within-population moments, five of which are for within the hub population and the remaining five for within the leaf population:

$$\frac{d\bar{S}_H}{dt} = \mu N - \frac{\beta}{N}(1-k\sigma)(C^H_{SI} + \bar{S}_H\bar{I}_H) - k\frac{\beta}{N}\sigma(\hat{C}_{S_H I_L} + \bar{S}_h\bar{I}_L) - \epsilon\bar{S}_H - \mu\bar{S}_H \tag{51}$$

$$\frac{d\bar{I}_H}{dt} = \frac{\beta}{N}(1-k\sigma)(C^H_{SI} + \bar{S}_H\bar{I}_H) + k\frac{\beta}{N}\sigma(\hat{C}_{S_H I_L} + \bar{S}_H\bar{I}_L) + \epsilon\bar{S}_H - \gamma\bar{I}_H - \mu\bar{I}_H \tag{52}$$

$$\begin{aligned}\frac{dC^H_{SS}}{dt} &= \mu N + \frac{\beta}{N}\left(1-\sigma\right)\bar{S}_H\bar{I}_L + \epsilon\bar{S}_H - \mu\bar{S}_H - 2\left(\frac{\beta}{N}(1-k\sigma)\bar{I}_H + k\frac{\beta}{N}\sigma\bar{I}_L + \epsilon + \mu\right)C^H_{SS} \\ &\quad - \frac{\beta}{N}(1-k\sigma)(2\bar{S}_H - 1)C^H_{SI} - k\frac{\beta}{N}\sigma(2\bar{S}_H - 1)\hat{C}_{S_H I_L}\end{aligned} \tag{53}$$

$$\begin{aligned}\frac{dC^H_{II}}{dt} &= \frac{\beta}{N}(1-k\sigma)\bar{S}_H\bar{I}_H + k\frac{\beta}{N}\sigma\bar{S}_H\bar{I}_L + \epsilon\bar{S}_H + (\gamma+\mu)\bar{I}_H + 2\left(\frac{\beta}{N}(1-k\sigma)\bar{S}_H - (\gamma+\mu)\right)C^H_{II} \\ &\quad + \left(\frac{\beta}{N}(1-k\sigma)(2\bar{I}_H + 1) + 2k\frac{\beta}{N}\sigma\bar{I}_L + 2\epsilon\right)C^H_{SI} + 2k\frac{\beta}{N}\sigma\bar{S}_H\hat{C}^H_{II} + k\frac{\beta}{N}\sigma\hat{C}_{S_H I_L}\end{aligned} \tag{54}$$

$$\begin{aligned}\frac{dC^H_{SI}}{dt} &= -\frac{\beta}{N}(1-k\sigma)\bar{S}_H\bar{I}_H - k\frac{\beta}{N}\sigma\bar{S}_H\bar{I}_L - \epsilon\bar{S}_H - \mu\bar{I}_H + \left(\frac{\beta}{N}(1-k\sigma)\bar{I}_H + k\frac{\beta}{N}\sigma\bar{I}_L + \epsilon\right)C^H_{SS} \\ &\quad - \frac{\beta}{N}(1-k\sigma)\bar{S}_H C^H_{II} + \left(\frac{\beta}{N}(1-k\sigma)(\bar{S}_H - \bar{I}_H - 1) - k\frac{\beta}{N}\sigma\bar{I}_L - \epsilon - \gamma - 2\mu\right)C^H_{SI} - k\frac{\beta}{N}\sigma\bar{S}_H\hat{C}^H_{II} + k\frac{\beta}{N}\sigma(\bar{S}_H - 1)\hat{C}_{S} \text{[illegible]}\end{aligned} \tag{55}$$

$$\frac{d\bar{S}_L}{dt} = \mu N - \frac{\beta}{N}\left(1-\sigma\right)\left(C^L_{SI} + \bar{S}_L\bar{I}_L\right) - \frac{\beta}{N}\sigma(\hat{C}_{S_L I_H} + \bar{S}_L\bar{I}_H) - \epsilon\bar{S}_L - \mu\bar{S}_L \tag{56}$$

$$\frac{d\bar{I}_L}{dt} = \frac{\beta}{N}\left(1-\sigma\right)\left(C^L_{SI} + \bar{S}_L\bar{I}_L\right) + \frac{\beta}{N}\sigma(\hat{C}_{S_L I_H} + \bar{S}_L\bar{I}_H) + \epsilon\bar{S}_L - \gamma\bar{I}_L - \mu\bar{I}_L \tag{57}$$

$$\begin{aligned}\frac{dC^L_{SS}}{dt} &= \mu N + \frac{\beta}{N}\left(1-\sigma\right)\bar{S}_L\bar{I}_L + \frac{\beta}{N}\sigma\bar{S}_L\bar{I}_H + \epsilon\bar{S}_H - \mu\bar{S}_H - 2\left(\frac{\beta}{N}\left(1-\sigma\right)\bar{I}_L + \frac{\beta}{N}\sigma\bar{I}_H + \epsilon + \mu\right)C^L_{SS} \\ &\quad - \frac{\beta}{N}\left(1-\sigma\right)(2\bar{S}_L - 1)C^L_{SI} - \frac{\beta}{N}\sigma(2\bar{S}_L - 1)\hat{C}_{S_L I_H}\end{aligned} \tag{58}$$

$$\begin{aligned}\frac{dC^L_{II}}{dt} &= \frac{\beta}{N}\left(1-\sigma\right)\bar{S}_L\bar{I}_L + \frac{\beta}{N}\sigma\bar{S}_L\bar{I}_H + \epsilon\bar{S}_L + (\gamma+\mu)\bar{I}_L + 2\left(\frac{\beta}{N}\left(1-\sigma\right)\bar{S}_L - (\gamma+\mu)\right)C^L_{II} \\ &\quad + \left(\frac{\beta}{N}\left(1-\sigma\right)(2\bar{I}_L + 1) + 2\frac{\beta}{N}\sigma\bar{I}_H + 2\epsilon\right)C^L_{SI} + 2\frac{\beta}{N}\sigma\bar{S}_L\hat{C}^H_{II} + \frac{\beta}{N}\sigma\hat{C}_{S_L I_H}\end{aligned} \tag{59}$$

$$\begin{aligned}\frac{dC^L_{SI}}{dt} &= -\frac{\beta}{N}\left(1-\sigma\right)\bar{S}_L\bar{I}_L - \frac{\beta}{N}\sigma\bar{S}_L\bar{I}_H - \epsilon\bar{S}_L - \mu\bar{I}_L + \left(\frac{\beta}{N}\left(1-\sigma\right)\bar{I}_L + \frac{\beta}{N}\sigma\bar{I}_H + \epsilon\right)C^L_{SS} \\ &\quad - \frac{\beta}{N}\left(1-\sigma\right)\bar{S}_L C^L_{II} + \left(\frac{\beta}{N}\left(1-\sigma\right)(\bar{S}_L - \bar{I}_L - 1) + \frac{\beta}{N}\sigma\bar{I}_H - \epsilon - \gamma - 2\mu\right)C^L_{SI} - \frac{\beta}{N}\sigma\bar{S}_L\hat{C}^H_{II} + k\frac{\beta}{N}\sigma(\bar{S}_L - 1)\hat{C}_{S_L H_H} \text{[illegible]}\end{aligned} \tag{60}$$

The remaining 7 equations are for the between-population moments:

$$\frac{d\hat{C}^H_{SS}}{dt} = \frac{\beta}{N}\sigma\bar{S}_L C^H_{SI} - \frac{\beta}{N}\sigma\bar{S}_H C^L_{SI} - \left(\frac{\beta}{N}(1-k\sigma)\bar{I}_H + k\frac{\beta}{N}\sigma\bar{I}_L + \frac{\beta}{N}\left(1-\sigma\right)\bar{I}_L + \frac{\beta}{N}\sigma\bar{I}_H + 2\epsilon + 2\mu\right)\hat{C}^H_{SS}$$
$$- \frac{\beta}{N}\left(1-\sigma\right)\bar{S}_L\hat{C}_{S_H I_L} - \frac{\beta}{N}(1-k\sigma)\bar{S}_H\hat{C}_{S_L I_H} - (k-1)\frac{\beta}{N}\sigma\bar{S}_H\hat{C}^L_{SI} \quad (61)$$

$$\frac{d\hat{C}^L_{SS}}{dt} = -2\left[\left(\frac{\beta}{N}\left(1-\sigma\right)\bar{I}_L + \frac{\beta}{N}\sigma\bar{I}_H + \epsilon + \mu\right)\hat{C}^L_{SS} + \frac{\beta}{N}\sigma\bar{S}_L\hat{C}_{S_L I_H} + \frac{\beta}{N}\left(1-\sigma\right)\bar{S}_L\hat{C}^L_{SI}\right] \quad (62)$$

$$\frac{d\hat{C}^H_{II}}{dt} = \frac{\beta}{N}\sigma\bar{S}_L C^H_{II} + \frac{\beta}{N}\sigma\bar{S}_H C^L_{II} + \left(\frac{\beta}{N}(1-k\sigma)\bar{S}_H + \frac{\beta}{N}\left(1-\sigma\right)\bar{S}_L - 2(\gamma+\mu)\right)\hat{C}^H_{II} + (k-1)\frac{\beta}{N}\sigma\bar{S}_H\hat{C}^L_{II}$$
$$+ \left(\frac{\beta}{N}(1-k\sigma)\bar{I}_H + k\frac{\beta}{N}\sigma\bar{I}_L + \epsilon\right)\hat{C}_{S_H I_L} + \left(\frac{\beta}{N}\left(1-\sigma\right)\bar{I}_L + \frac{\beta}{N}\sigma\bar{I}_H + \epsilon\right)\hat{C}_{S_L I_H} \quad (63)$$

$$\frac{d\hat{C}^L_{II}}{dt} = 2\left[\frac{\beta}{N}\sigma\bar{S}_L\hat{C}^H_{II} + \left(\frac{\beta}{N}\left(1-\sigma\right) - (\gamma+\mu)\right)\hat{C}^L_{II} + \left(\frac{\beta}{N}\left(1-\sigma\right)\bar{I}_L + \frac{\beta}{N}\sigma\bar{I}_H\epsilon\right)\hat{C}^L_{SI}\right] \quad (64)$$

$$\frac{d\hat{C}_{S_H I_L}}{dt} = -\frac{\beta}{N}\sigma\bar{S}_H C^L_{II} + \frac{\beta}{N}\sigma\bar{S}_L\bar{S}I^H + \left(\frac{\beta}{N}\left(1-\sigma\right)\bar{I}_L + \frac{\beta}{N}\sigma\bar{I}_H + \epsilon\right)\hat{C}^H_{SS} - \frac{\beta}{N}(1-k\sigma)\bar{S}_H\hat{C}^H_{II}$$
$$- (k-1)\frac{\beta}{N}\sigma\bar{S}_H\hat{C}^L_{II} + \left(\frac{\beta}{N}\left(1-\sigma\right)\bar{S}_L - \frac{\beta}{N}(1-k\sigma)\bar{I}_H - k\frac{\beta}{N}\sigma\bar{I}_L - \epsilon - \gamma - 2\mu\right)\hat{C}_{S_H I_L} \quad (65)$$

$$\frac{d\hat{C}_{S_L I_H}}{dt} = -\frac{\beta}{N}\sigma\bar{S}_L C^H_{II} + \frac{\beta}{N}\sigma\bar{S}_H C^L_{SI} + \left(\frac{\beta}{N}(1-k\sigma)\bar{I}_H + k\frac{\beta}{N}\sigma\bar{I}_L + \epsilon\right)\hat{C}^H_{SS} - \frac{\beta}{N}\left(1-\sigma\right)\bar{S}_L\hat{C}^H_{II}$$
$$+ \left(\frac{\beta}{N}(1-k\sigma)\bar{S}_H - \frac{\beta}{N}\left(1-\sigma\right)\bar{I}_L - \frac{\beta}{N}\sigma\bar{I}_H - \epsilon - \gamma - 2\mu\right)\hat{C}_{S_L I_H} + (k-1)\frac{\beta}{N}\sigma\bar{S}_H\hat{C}^L_{SI} \quad (66)$$

$$\frac{d\hat{C}^L_{SI}}{dt} = \left(\frac{\beta}{N}\left(1-\sigma\right)\bar{I}_L + \frac{\beta}{N}\sigma\bar{I}_H + \epsilon\right)\hat{C}^L_{SS} - \frac{\beta}{N}\sigma\bar{S}_L\hat{C}^H_{II} - \frac{\beta}{N}\left(1-\sigma\right)\bar{S}_L\hat{C}^L_{II} + \frac{\beta}{N}\sigma\bar{S}_L\hat{C}_{S_L I_H}$$
$$+ \left(\frac{\beta}{N}\left(1-\sigma\right)(\hat{S}_L - \hat{I}_L) - \frac{\beta}{N}\sigma\bar{I}_H - \epsilon - \gamma - 2\mu\right)\hat{C}^L_{SI} \quad (67)$$

### 8 Derivation of the approximation for the star network

For the star network with $P$ subpopulations on the star network, where the coupling between interacting populations is $\sigma \in [0, 1/k], k = P-1$, we can show that the correlation between the number of infected individuals in the hub population and a leaf population, $\rho_H$, and the correlation between the number of infected individuals in two leaf populations, $\rho_L$ are solution to the following pair of simultaneous equations:

$$\rho_H = \sqrt{\frac{C_{II}^{H*}}{C_{II}^{L*}}} \frac{\sigma}{\frac{\bar{S}_H^*}{\bar{S}_L^*}(\xi_H + k\sigma) + \xi_L + \sigma} + \sqrt{\frac{C_{II}^{L*}}{C_{II}^{H*}}} \frac{\sigma}{\xi_H + k\sigma + \frac{\bar{S}_L^*}{\bar{S}_H^*}(\xi_L + \sigma)} (1 - (1-k)\rho_L) - \Delta_H \tag{68}$$

$$\rho_L = \sqrt{\frac{C_{II}^{H*}}{C_{II}^{L*}}} \frac{\sigma}{\xi_L + \sigma} \rho_H + \Delta_L, \tag{69}$$

where

$$\xi_H = \frac{N(\gamma + \mu) - \beta \bar{S}_H^*}{\beta \bar{S}_H^*}, \tag{70}$$

$$\xi_L = \frac{N(\gamma + \mu) - \beta \bar{S}_L^*}{\beta \bar{S}_L^*} \tag{71}$$

and

$$\Delta_H = \frac{\beta(1-k\sigma)\bar{I}_H^* + k\beta\sigma \bar{I}_L^* + N\epsilon}{2N(\gamma+\mu) - \beta(1-k\sigma)\bar{S}_H^* - \beta(1-\sigma)\bar{S}_L^*} \frac{\hat{C}_{S_H I_L}^*}{\sqrt{C_{II}^{H*} C_{II}^{L*}}} + \frac{\beta(1-\sigma)\bar{I}_L^* + \beta\sigma \bar{I}_H^* + N\epsilon}{2N(\gamma+\mu) - \beta(1-k\sigma)\bar{S}_H^* - \beta(1-\sigma)\bar{S}_L^*} \frac{\hat{C}_{S_L I_H}^*}{\sqrt{C_{II}^{H*} C_{II}^{L*}}} \tag{72}$$

$$\Delta_L = \frac{\beta(1-\sigma)\bar{I}_L^* + \beta\sigma \bar{I}_H^* + N\epsilon}{N(\gamma+\mu) - \beta(1-\sigma)\bar{S}_L^*} \frac{\hat{C}_{SI}^{L*}}{C_{II}^{L}}. \tag{73}$$

We derive Equation (68) using the moment equation for the covariance between the number of infected individuals in the hub and a leaf population, $\hat{C}_{II}^H$:

$$\frac{d\hat{C}_{II}^H}{dt} = \frac{\beta}{N}\sigma \bar{S}_L C_{II}^H + \frac{\beta}{N}\sigma \bar{S}_H C_{II}^L + \left(\frac{\beta}{N}(1-k\sigma)\bar{S}_H + \frac{\beta}{N}(1-\sigma)\bar{S}_L - 2(\gamma+\mu)\right)\hat{C}_{II}^H + (k-1)\frac{\beta}{N}\sigma \bar{S}_H \hat{C}_{II}^L + \left(\frac{\beta}{N}(1-k\sigma)\bar{I}_H + \frac{\beta}{N}k\sigma \bar{I}_L + \epsilon\right)\hat{C}_{S_H I_L} + \left(\frac{\beta}{N}(1-\sigma)\bar{I}_L + \frac{\beta}{N}\sigma \bar{I}_H + \epsilon\right)\hat{C}_{S_L I_H}. \tag{74}$$

At equilibrium $d\hat{C}_{II}^H/dt = 0$, and if we divide by $\sqrt{C_{II}^{H*} C_{II}^{L*}}/N$ then

$$0 = \sqrt{\frac{C_{II}^{H*}}{C_{II}^{L*}}} \beta\sigma \bar{S}_L^* + \sqrt{\frac{C_{II}^{L*}}{C_{II}^{H*}}} \beta\sigma \bar{S}_H^* + \left(\beta(1-k\sigma)\bar{S}_H^* + \beta(1-\sigma)\bar{S}_L^* - 2N(\gamma+\mu)\right)\rho_H + \sqrt{\frac{C_{II}^{L*}}{C_{II}^{H*}}}(k-1)\beta\sigma \bar{S}_H^* \rho_L + \left(\beta(1-k\sigma)\bar{I}_H^* + \beta k\sigma \bar{I}_L^* + N\epsilon\right)\frac{\hat{C}_{S_H I_L}^*}{\sqrt{C_{II}^{H*} C_{II}^{L*}}} + \left(\beta(1-\sigma)\bar{I}_L^* + \beta\sigma \bar{I}_H^* + N\epsilon\right)\frac{\hat{C}_{S_L I_H}^*}{\sqrt{C_{II}^{H*} C_{II}^{L*}}}, \tag{75}$$

and hence we have the following approximation for the correlation between the number of infected individuals in the hub and a leaf population:

$$\begin{aligned}
\rho_H &= \sqrt{\frac{C_{II}^{H*}}{C_{II}^{L*}}} \frac{\beta\sigma\bar{S}_L^*}{2N(\gamma+\mu) - \beta(1-k\sigma)\bar{S}_H^* - \beta(1-\sigma)\bar{S}_L^*} \\
&\quad + \sqrt{\frac{C_{II}^{L*}}{C_{II}^{H*}}} \frac{\beta\sigma\bar{S}_H^*}{2N(\gamma+\mu) - \beta(1-k\sigma)\bar{S}_H^* - \beta(1-\sigma)\bar{S}_L^*}(1-(1-k)\rho_L) - \Delta_H \\
&= \sqrt{\frac{C_{II}^{H*}}{C_{II}^{L*}}} \frac{\sigma}{\frac{N(\gamma+\mu)-\beta(1-k\sigma)\bar{S}_H^*}{\beta\bar{S}_L^*} + \frac{N(\gamma+\mu)-\beta(1-\sigma)\bar{S}_L^*}{\beta\bar{S}_L^*}} \\
&\quad + \sqrt{\frac{C_{II}^{L*}}{C_{II}^{H*}}} \frac{\sigma}{\frac{N(\gamma+\mu)-\beta(1-k\sigma)\bar{S}_H^*}{\beta\bar{S}_H^*} + \frac{N(\gamma+\mu)-\beta(1-\sigma)\bar{S}_L^*}{\beta\bar{S}_H^*}}(1-(1-k)\rho_L) - \Delta_H \\
&= \sqrt{\frac{C_{II}^{H*}}{C_{II}^{L*}}} \frac{\sigma}{\frac{\bar{S}_H^*}{\bar{S}_L^*}\left(\frac{N(\gamma+\mu)-\beta\bar{S}_H^*}{\beta\bar{S}_H^*} + k\sigma\right) + \frac{N(\gamma+\mu)-\beta\bar{S}_L^*}{\beta\bar{S}_L^*} + \sigma} \\
&\quad + \sqrt{\frac{C_{II}^{L*}}{C_{II}^{H*}}} \frac{\sigma}{\frac{N(\gamma+\mu)-\beta\bar{S}_H^*}{\beta\bar{S}_H^*} + k\sigma + \frac{\bar{S}_L^*}{\bar{S}_H^*}\left(\frac{N(\gamma+\mu)-\beta\bar{S}_L^*}{\beta\bar{S}_L^*} + \sigma\right)}(1-(1-k)\rho_L) - \Delta_H \\
&= \sqrt{\frac{C_{II}^{H*}}{C_{II}^{L*}}} \frac{\sigma}{\frac{\bar{S}_H^*}{\bar{S}_L^*}(\xi_H + k\sigma) + \xi_L + \sigma} + \sqrt{\frac{C_{II}^{L*}}{C_{II}^{H*}}} \frac{\sigma}{\xi_H + k\sigma + \frac{\bar{S}_L^*}{\bar{S}_H^*}(\xi_L+\sigma)}(1-(1-k)\rho_L) - \Delta_H. \qquad (76)
\end{aligned}$$

Similarly, we derive Equation (69) using the moment equations for the covariance between the number of infected individuals in two distinct leaf populations, $\hat{C}_{II}^L$:

$$\frac{d\hat{C}_{II}^L}{dt} = 2\left[\frac{\beta}{N}\sigma\bar{S}_L\hat{C}_{II}^H + \left(\frac{\beta}{N}(1-\sigma)\bar{S}_L - \gamma - \mu\right)\hat{C}_{II}^L + \left(\frac{\beta}{N}(1-\sigma)\bar{I}_L + \frac{\beta}{N}\sigma\bar{I}_H + \epsilon\right)\hat{C}_{SI}^L\right]. \qquad (77)$$

At equilibrium $d\hat{C}_{II}^L/dt = 0$ and if we divide by $2C_{II}^{L*}/N$, then

$$0 = \sqrt{\frac{C_{II}^{H*}}{C_{II}^{L*}}}\beta\sigma\bar{S}_L^*\rho_H + \left(\beta(1-\sigma)\bar{S}_L^* - N(\gamma+\mu)\right)\rho_L + \left(\beta(1-\sigma)\bar{I}_L^* + \beta\sigma\bar{I}_H^* + N\epsilon\right)\frac{\hat{C}_{SI}^{L*}}{C_{II}^{L*}}, \qquad (78)$$

and hence we have the following approximation for the correlation between the number of infected individuals in two leaf populations:

$$\begin{aligned}
\rho_L &= \sqrt{\frac{C_{II}^{H*}}{C_{II}^{L*}}} \frac{\beta\sigma\bar{S}_L^*}{N(\gamma+\mu) - \beta(1-\sigma)\bar{S}_L^*}\rho_H + \frac{\beta(1-\sigma)\bar{I}_L^* + \beta\sigma\bar{I}_H^* + N\epsilon}{N(\gamma+\mu) - \beta(1-\sigma)\bar{S}_L^*}\frac{\hat{C}_{SI}^{L*}}{C_{II}^{L*}} \\
&= \sqrt{\frac{C_{II}^{H*}}{C_{II}^{L*}}} \frac{\sigma}{\frac{N(\gamma+\mu)-\beta\bar{S}_L^*}{\beta\bar{S}_L^*} + \sigma}\rho_H + \Delta_L \\
&= \sqrt{\frac{C_{II}^{H*}}{C_{II}^{L*}}} \frac{\sigma}{\xi_L + \sigma}\rho_H + \Delta_L. \qquad (79)
\end{aligned}$$

#### 9 Correlations in the $D$-truncated $k$-regular tree network

In our main analysis we perform stochastic simulations $D$-truncated $k$-regular tree network. The correlation between the number of infected individuals in subpopulations distance $d \ll D$ apart is calculated between the origin subpopulation and another subpopulation distance $d$ away. If $D$ is sufficiently large, then these correlations will be approximately the same as in the full $k$-regular tree network. We show that the threshold for $D$ is a function of both the coupling $\sigma$ and the number of neighbours $k$.

We calculate the correlation $\rho_1$ between adjacent populations in the $D$-truncated $k$-regular tree network for $k = 2, 4$ and a range of values for $D$ (Figure S2). For low coupling ($\sigma = 0.001, 0.01$), increasing $D$ has little effect on the correlation: this suggests that when coupling is low it is sufficient to take $D = 3$. However, for larger coupling ($\sigma = 0.1$) then as $D$ increases then the correlation initially decreases, and then plateaus: for both $k = 2$ and $k = 4$ the correlation plateaus around $D = 5$.

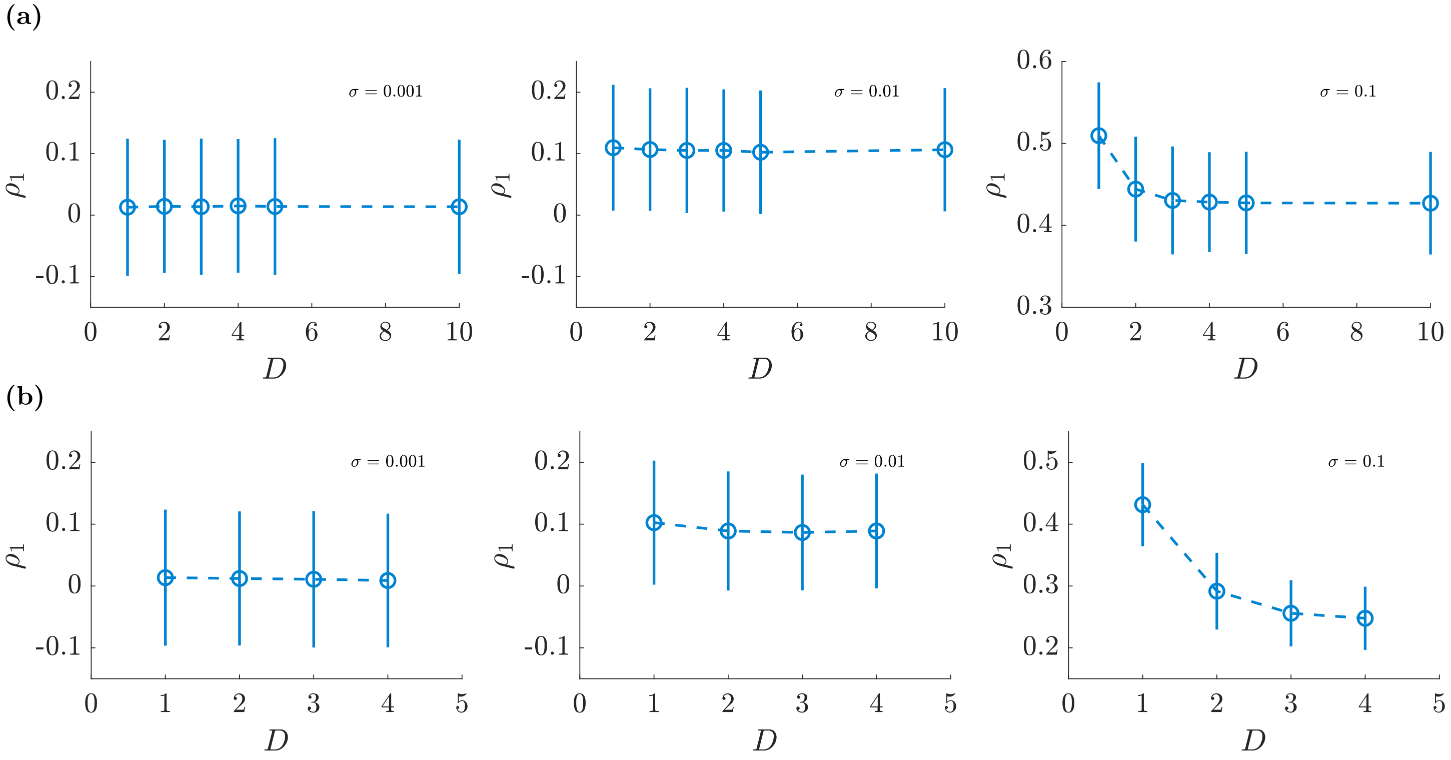


**Figure S2.** The correlation $\rho_1$ between adjacent subpopulations in the $D$-truncated $k$-regular tree network for $\sigma = 0.001, 0.01, 0.1$, and (a) $k = 2$, and (b) $k = 4$. Parameter values represent a measles-like endemic disease in the UK ($N = 10^5, \mu = 5.5 \times 10^{-5}, R_0 = 17, \gamma^{-1} = 13$ and $\epsilon = 5.5 \times 10^{-5}$). We generate 1000 realisations of the process for each value of $\sigma$ and calculate the correlation as a time-weighted Pearson correlation coefficient for $50 \leq t \leq 200$; error bars represent $\pm 2$ standard deviations.
